# Endocytic vesicles act as vehicles for glucose uptake in response to growth factor stimulation

**DOI:** 10.1101/2023.07.23.550235

**Authors:** Ryouhei Tsutsumi, Beatrix Ueberheide, Feng-Xia Liang, Benjamin G. Neel, Ryuichi Sakai, Yoshiro Saito

## Abstract

Glycolysis is a fundamental cellular process, yet its regulatory mechanisms remain incompletely understood. Here, we show that a subset of glucose transporter 1 (GLUT1/SLC2A1) co-endocytoses with platelet-derived growth factor (PDGF) receptor (PDGFR) upon PDGF-stimulation. Furthermore, multiple glycolytic enzymes localize to these endocytosed PDGFR/GLUT1-containing vesicles adjacent to mitochondria. Contrary to current models, which emphasize the importance of glucose transporters on the cell surface, we find that PDGF-stimulated glucose uptake depends on receptor/transporter endocytosis. Our results suggest that growth factors generate glucose-loaded endocytic vesicles that deliver glucose to the glycolytic machinery in proximity to mitochondria, and argue for a new layer of regulation for glycolytic control governed by cellular membrane dynamics.

## Introduction

High rates of glycolysis correlate with cell stemness and cancer (1, 2), yet the underlying mechanisms are incompletely understood. The rate-determining steps of glycolysis include uptake of extracellular glucose by transporters and subsequent phosphorylation by hexokinases (HKs) to prevent re-release of glucose to the extracellular space. Among the mammalian facilitated glucose transporter (GLUT) family members SLC2A1-14, the ubiquitously expressed GLUT1 (SLC2A1), the renal tubular cell-, hepatocyte-and pancreatic β cell-specific GLUT2 (SLC2A2), the neuron-and placenta-specific GLUT3 (SLC2A3), and the adipose tissue-and striated muscle-specific GLUT4 (SLC2A4) are well characterized (3). GLUT4 is the only glucose transporter known to dynamically increase glucose uptake without the need for new transcription/translation; in response to insulin, it translocates from intracellular structures to the plasma membrane (3, 4). However, earlier work showed that stimulation of fibroblasts, which do not express GLUT4, with epidermal growth factor (EGF) resulted in elevation of glucose uptake within 15 min (5). This observation cannot be explained by current models for glycolysis regulation in such cells, which feature transcriptional upregulation of glucose transporters and/or glycolytic enzymes (1). One study reported that fibroblasts treated with 12-*O*-tetradecanoylphorbol-13-acetate (TPA) activate GLUT1 via PKC-dependent phosphorylation on S226 (6), although whether PKC-dependent phosphorylation is fully responsible for promoting growth factor-evoked glucose uptake has remained unclear.

Growth factors bind and activate their cognate receptor tyrosine kinases (RTKs) at the plasma membrane, resulting in protein tyrosyl phosphorylation-dependent signaling and receptor endocytosis (7, 8). Endocytosis results in RTK inactivation/degradation unless the RTK is recycled to the plasma membrane. However, multiple studies also implicate intracellular vesicles containing active RTKs as signaling platforms (9, 10). We developed a method, analogous to one reported previously (11), that enables efficient and quick recovery of growth factor-evoked, RTK-containing endocytic vesicles. Here, we report that upon growth factor stimulation, GLUT1 and glycolytic enzymes co-localize with such vesicles. Furthermore, our data suggest that receptor/GLUT1 “co-endocytosis” serves to deliver glucose to glycolytic enzymes in proximity to mitochondria, allowing rapid adaptation of cellular glucose metabolism to mitogenic signals.

## Results

### Nanoparticle-based isolation identified PDGFR-containing endocytic vesicle-associated proteins

PDGFR has α and β isoforms, which homo-or heterodimerize in response to different dimeric PDGF isoforms (PDGF-AA, PDGF-BB, PDGF-AB, PDGF-CC, PDGF-DD) (12). Serum-starved Swiss 3T3 mouse fibroblasts, which express PDGFRα and β, were treated for 5 min with biotinylated PDGF-BB bound to streptavidin-coated magnetic iron oxide nanoparticles with a diameter of 10 nm. These particles bind to surface PDGFRs and, following PDGF stimulation, are transported with the receptor into endocytic vesicles (∼100 nm) (Fig. S1A, B). Nanoparticles remaining on the cell surface were stripped by acid washes, and post-nuclear supernatants were passed through a magnetic separation column to enrich for nanoparticle-containing structures (Fig. 1A), resolved by SDS-PAGE, and silver-stained (Fig. 1B). Treating cells with the PDGF-BB-conjugated nanoparticles at 4°C to inhibit receptor endocytosis, or treatment with nanoparticles and unconjugated PDGF-BB at 37°C, resulted in recovery of significantly fewer proteins (Fig. 1B, S1C).

**Figure 1.**
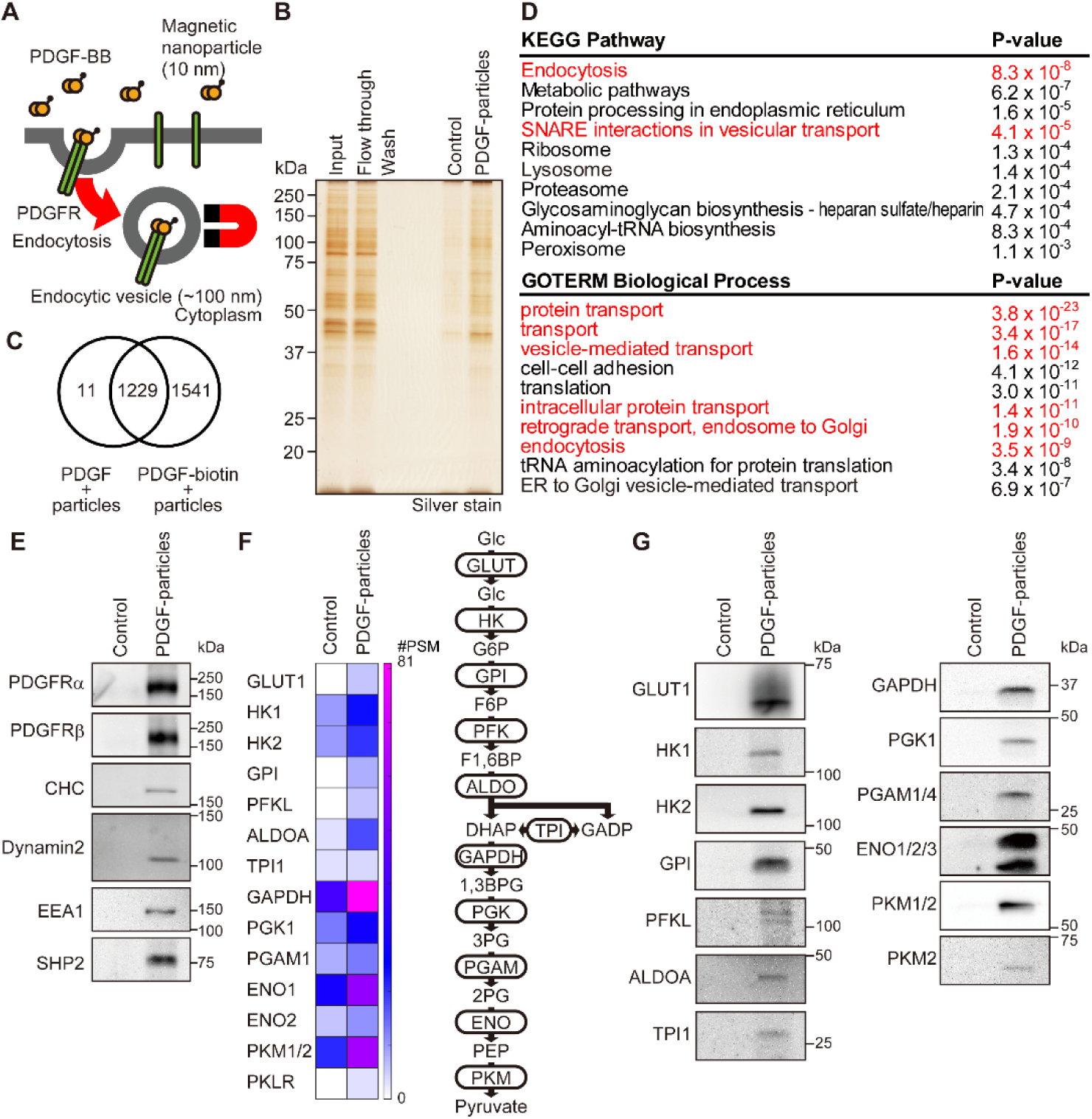
Isolation of PDGFR endocytic vesicles utilizing magnetic nanoparticles. (A) Schematic showing strategy for endocytic vesicle isolation. (B) Fractions magnetically isolated from post-nuclear supernatants of PDGF-BB-biotin-conjugated nanoparticle-treated (PDGF-particle) or unconjugated PDGF-BB plus nanoparticle-treated (control) Swiss 3T3 fibroblasts were analyzed by SDS-PAGE and silver staining. Post-nuclear supernatant (input), flowthrough, and wash fractions were from PDGF-BB-biotin-conjugated nanoparticle-treated cells. A representative image from one of 5 independent experiments is shown. (C) Venn diagram shows numbers of proteins identified in the control and endocytic vesicle fractions by LC-MS/MS, as detailed in Table S1. Data are from a single experiment. (D) Gene ontology (GO) enrichment analyses of proteins uniquely identified in the endocytic vesicle fraction. Annotations related to endocytosis or vesicle transport are in red. (E) Fractions magnetically isolated from post-nuclear supernatants of PDGF-BB-biotin-conjugated nanoparticle-treated (PDGF-particle) or unconjugated PDGF-BB plus nanoparticle-treated (control) Swiss 3T3 fibroblasts were subjected to immunoblotting with the indicated antibodies. Representative images are shown from one of 2 independent experiments. (F) Heatmap representing numbers of peptide spectrum matches (#PSMs) of glycolysis-related proteins obtained by LC-MS/MS. A schematic of glycolysis is shown on the right. (G) Fractions magnetically isolated from post-nuclear supernatants of PDGF-BB-biotin-conjugated nanoparticle-treated (PDGF-particle) or unconjugated PDGF-BB plus nanoparticle-treated (control) Swiss 3T3 fibroblasts were subjected by immunoblotting with indicated antibodies. Representative images are shown from one of 2 independent experiments.

Liquid chromatography-tandem mass spectrometric (LC-MS/MS) analysis of isolates from cells stimulated with PDGF-BB plus unconjugated nanoparticles (control) or with PDGF-BB-conjugated nanoparticles (endocytic vesicle fraction) identified 2,781 proteins at a threshold of 2 unique peptides per protein. Of these, 1,541 were unique to the endocytic vesicle fraction (Fig. 1C, Table S1) and were enriched for proteins annotated as having endocytosis-or vesicular transport-related functions (Fig. 1D), supporting the validity of our method. The proteins detected in response to either type of stimulation could reflect ligand-stimulated bulk endocytic transport of nanoparticles into cells as reported previously (13). Even so, we detected more of these proteins, especially endocytosis-related proteins, in the PDGF-BB-conjugated nanoparticle fraction than in the unconjugated PDGF-BB plus nanoparticle control (Fig. S1D). Immunoblotting experiments confirmed that PDGFR and endocytosis-related proteins, including clathrin heavy chain (CHC), dynamin 2, and early endosome antigen 1 (EEA1), were detected almost exclusively in the endocytic vesicle fraction (Fig. 1E). The endocytic vesicle fraction also contained proteins related to vesicle maturation and transportation, as well as cell signaling (Table S1, Fig. S1E, F), comporting with the notion that these vesicles function as signaling platforms (9, 10). In addition, this fraction contained Golgi apparatus, endoplasmic reticulum (ER) and mitochondrial proteins (Fig. S1E, Table S1). These could represent organellar contamination but also might indicate recovery of organelle-endocytic vesicle contacts.

### PDGFR co-endocytoses with GLUT1 and colocalizes with most glycolytic enzymes

Unexpectedly, GLUT1, as well as most glycolytic enzymes, including hexokinase (HK1 and HK2), glucose-6-phosphate isomerase (GPI), phosphofructokinase (PFKL), aldolase (ALDOA), triosephosphate isomerase (TPI1), glyceraldehyde 3-phosphate dehydrogenase (GAPDH), phosphoglycerate kinase (PGK1), phosphoglycerate mutase (PGAM1), enolase (ENO1/2), and pyruvate kinase (PKM1/2, PKLR), also were enriched in the endocytic vesicle fraction by MS (Fig. 1F) and by immunoblotting (Fig. 1G). In parallel (and to rule out their artifactual recovery owing to organellar contamination of the vesicle fraction), we monitored the subcellular localization of these proteins by confocal immunofluorescence microscopy. As expected, in PDGF-stimulated Swiss 3T3 fibroblasts, PDGFRα and β colocalized in vesicular structures, which also stained with the early endosome marker EEA1 (Fig. S2A). Immunostaining of serum-starved fibroblasts revealed GLUT1 localization at the plasma membrane and in perinuclear punctate structures, whereas stimulation with 50 ng/ml PDGF-BB for 10 min resulted in relocation of GLUT1 to cytoplasmic vesicular structures that merged with PDGFRα (Fig. 2A, S2B, S3A, S3B). Colocalization was also observed when cell surface GLUT1 was pre-labeled before PDGF stimulation by capitalizing on recombinant version of the GLUT1-binding region of human T cell leukemia virus envelope glycoprotein (14) (Fig. 2B). In concert, these data show that GLUT1 is co-endocytosed with PDGFRα/β following ligand stimulation.

**Figure 2.**
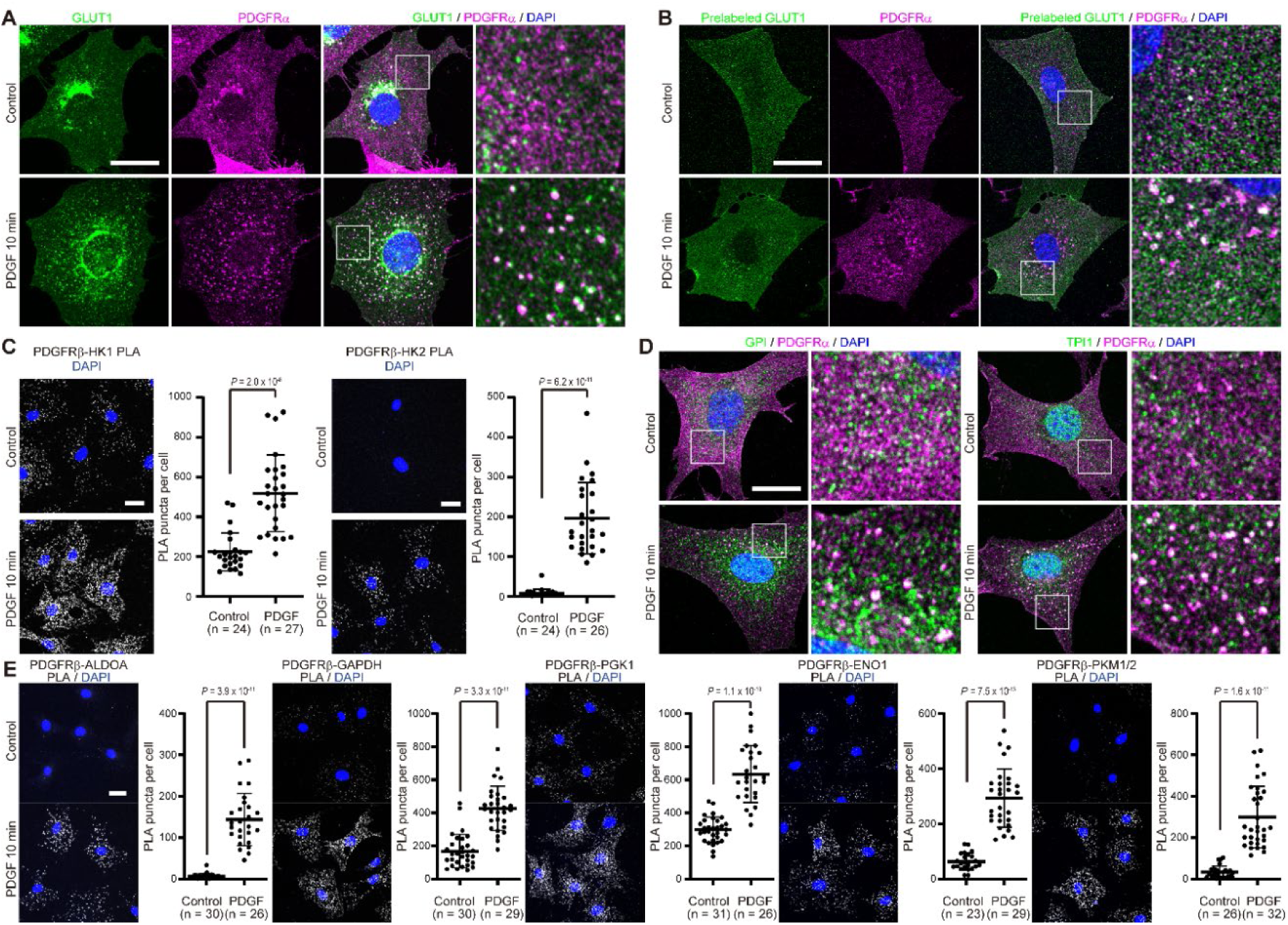
GLUT1 and glycolytic enzymes localize to PDGFR endocytic vesicles. (A) Serum-starved Swiss 3T3 fibroblasts were stimulated with PDGF-BB (50 ng/ml) for 10 min or left unstimulated (control), and then immunostained with anti-GLUT1 (green) and anti-PDGFRα (magenta) antibodies. Nuclei were stained with DAPI (blue). Higher magnification images of the boxed regions are shown on the right. Representative images are shown for each condition from one of 2 independent experiments. (B) Cell surface GLUT1 in serum-starved Swiss 3T3 cells was labeled with recombinant GLUT1-binding region of human T cell leukemia virus envelope glycoprotein for 20 min on ice, and then the cells were stimulated with PDGF-BB (50 ng/ml) for 10 min at 37°C or left unstimulated and immunostained for surface and internalized GLUT1 (green) and PDGFRα (magenta). Nuclei were stained with DAPI (blue). Higher magnification images of the boxed regions are shown on the right. Representative images are shown for each condition from one of 2 independent experiments. (C) Serum-starved Swiss 3T3 fibroblasts were stimulated with PDGF-BB (50 ng/ml) for 10 min or left unstimulated, and then subjected to PLA with anti-PDGFRβ and anti-HK1 or -HK2 antibodies (gray). Nuclei were stained with DAPI (blue). PLA signals in the indicated number of cells were counted and plotted in the graphs. Bars represent mean ± SD of PLA signals per cell. P values were calculated using two-tailed Welch’s *t* test. Representative data are shown from one of 2 independent experiments. (D) Serum-starved and PDGF-BB-stimulated or unstimulated cells (as above) were immunostained with anti-GPI or anti-TPI (green) and anti-PDGFRα (magenta) antibodies. Higher magnification images of the boxed regions are shown on the right. Nuclei were stained with DAPI (blue). Representative images are shown for each condition from one of 2 independent experiments. (E) Serum-starved and PDGF-BB stimulated or unstimulated cells were subjected to PLA with anti-PDGFRβ and the indicated antibodies. PLA signals were counted and plotted in the graphs. Bars represent mean ± SD of PLA signals per cell. P values were calculated using two-tailed Welch’s *t* test. Representative data are shown from one of 2 independent experiments. Scale bars: 20 μm.

HK1 and HK2 catalyze the first reaction of glycolysis, phosphorylation at the position 6 hydroxyl group of glucose. Both localize to the cytoplasmic surface of the mitochondrial outer membrane (Fig. S2C, S3A, S3B), although a fraction of HK2 also is reported to reside in the cytoplasm (15). Co-staining of PDGFRα and HK1 or HK2 in PDGF-stimulated fibroblasts did not reveal ligand-dependent relocation of HKs (Fig. S2C); instead, we observed that PDGFRα-containing vesicles localized in proximity to mitochondria. Recent studies have reported inter-organelle interactions between endosomes/lysosomes and mitochondria through RAB7, MFN2, VPS13, and/or VDAC (16–19), which were also enriched in our endocytic vesicle fractionation (Fig S1E, Table S1). We therefore asked whether endocytic vesicles and mitochondria interact after PDGF-stimulation by employing proximity ligation assay (PLA), which detects different antibodies that reside within 40 nm of each other (20). PLA signals between PDGFRβ and HK1 or HK2, respectively, were enhanced significantly following PDGF-stimulation (Fig. 2C, S3C), indicating that PDGFR-containing endocytic vesicles are transported to within nanometers of mitochondria. Importantly, PLA signals were abolished in cells treated with Hk1 or Hk2 siRNAs, respectively (Fig. S3C).

GPI and TPI were observed predominantly in the cytoplasm in serum-starved cells, but PDGF-stimulation resulted in their relocation to PDGFRα-containing vesicles (Fig. 2D, S3A, S3B). Other glycolytic enzymes, including ALDOA, GAPDH, PGK1, ENO1, and PKM1/2 showed strong cytoplasmic staining regardless of growth factor stimulation, but also partial co-localization to the periphery of PDGFR-containing vesicles (Fig. S2D, S3A, S3B) in PDGF-stimulated cells. PDGF-stimulation significantly increased PLA signals between PDGFRβ and these enzymes (Fig. 2E, S3C). Importantly, PDGF stimulation of serum-starved fibroblasts did not increase the levels of the glycolytic enzymes (Fig. S2E). We could not evaluate the localization of PFKL, PGAM1, or PKLR owing to a lack of antibodies available for immunostaining. Collectively, however, our results show that GLUT1 is co-endocytosed with PDGFR upon PDGF stimulation and suggest that these endocytic vesicles localize close to mitochondrial hexokinases and most, if not all, other glycolytic enzymes.

### PDGF-evoked GLUT1-mediated cellular glucose uptake is dependent on receptor endocytosis

Cell surface expression of glucose transporters is thought to determine the rate of extracellular glucose uptake. Accordingly, PDGF-induced GLUT1 internalization as described might be expected to decrease cellular glucose uptake (and thereby glycolysis). Yet it has long been known that growth factor stimulation rapidly augments glucose uptake and glycolysis (5, 21, 22). To explore this apparent paradox, we tested glucose uptake by Swiss 3T3 fibroblasts by incubating them in the presence of the non-metabolizable analog 2-deoxyglucose (2DG) and measuring the accumulation of 2-deoxyglucose-6-phosphate (2DG6P). Indeed, PDGF stimulation for 10 min nearly doubled glucose uptake (Fig. S4A). Basal and stimulation-evoked elevation of glucose uptake in Swiss 3T3 fibroblasts were dependent on GLUT1 because treatment with 50 nM BAY876, a specific GLUT1 inhibitor at this concentration (23), almost completely abolished 2DG6P formation without affecting PDGFR tyrosyl phosphorylation (Figs. 3A, S4B). On the other hand, surface and total GLUT1 levels decreased slightly after PDGF stimulation (Figs. S4C, D), indicating that PDGF-dependent upregulation of glucose uptake is not due to a rapid increase in surface expression of the transporter. We conclude that PDGF elicits glucose uptake despite endocytosis of the major glucose transporter GLUT1 in 3T3 fibroblasts.

**Figure 3.**
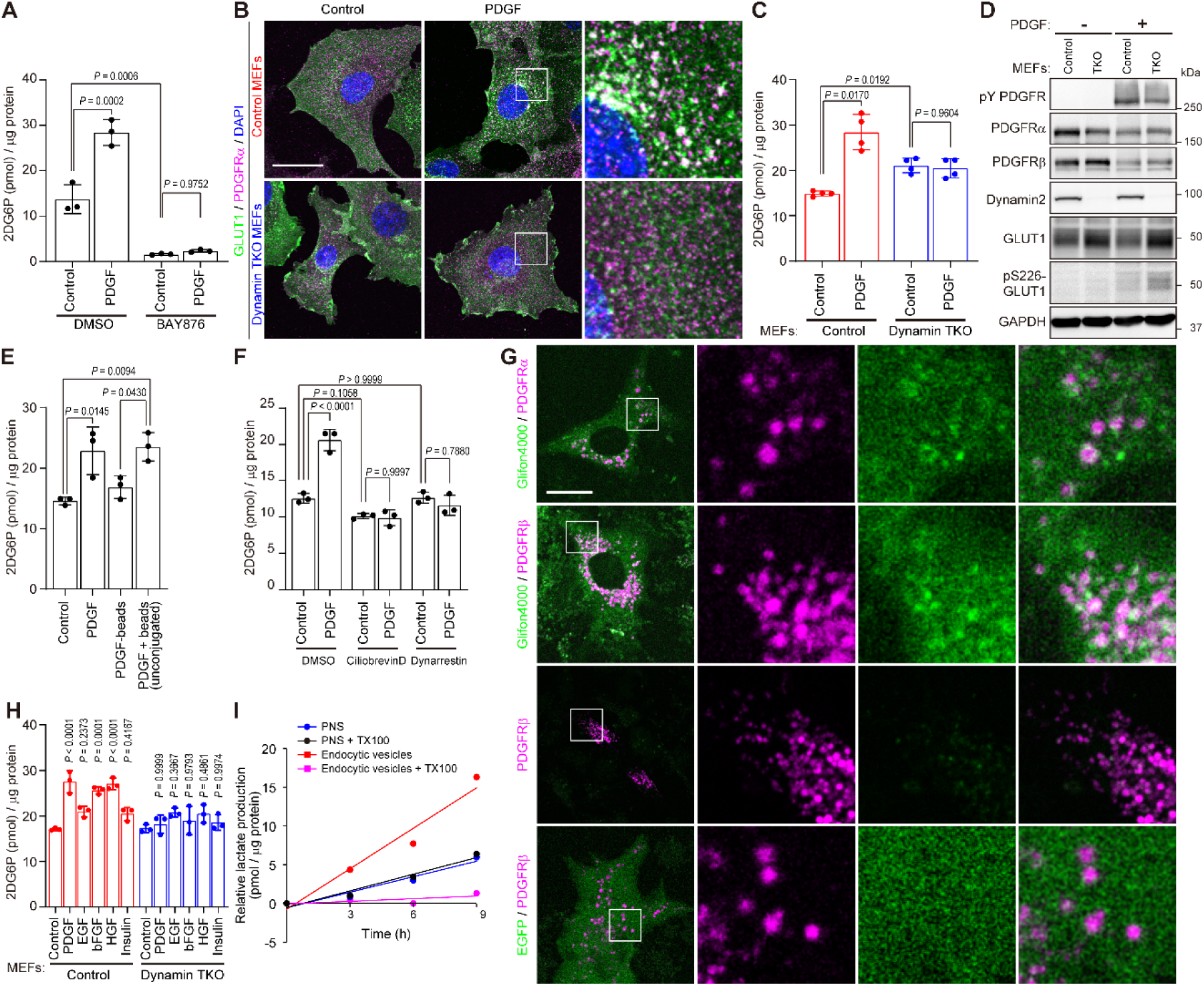
Growth factor-evoked glucose uptake requires receptor/GLUT1 endocytosis. (A) Serum-starved Swiss 3T3 fibroblasts were pre-treated with or without GLUT1-specific inhibitor BAY876 (50 nM) for 20 min and then stimulated with PDGF-BB (50 ng/ml) or left unstimulated (Control) in the presence of 2DG for 10 min. Graph shows average of 2DG6P in 3 wells for each condition (technical replicates, n = 3), normalized to total cellular protein. Error bars represent ± SD. P values were calculated using ANOVA and post-hoc Tukey’s tests. Representative data are shown from one of 2 independent experiments. (B) Conditional dynamin TKO MEFs were treated with 4-OHT (dynamin TKO MEFs) or left untreated (control MEFs), serum-starved, stimulated with 50 ng/ml PDGF-BB for 10 min, and immunostained with anti-GLUT1 (green) and anti-PDGFRα (magenta) antibodies. Higher magnification images of the boxed regions are shown on the right. Nuclei were stained with DAPI (blue). Representative data are shown from one of 2 independent experiments. Scale bar: 20 μm. (C) Serum-starved control or dynamin TKO MEFs were subjected to glucose uptake assay with or without PDGF-BB (50 ng/ml) in the presence of 2DG for 10 min. Graph shows average 2DG6P levels normalized to total cellular protein from 4 independent biological replicates (n = 4). Error bars represent ± SD. P values were calculated using RM one-way ANOVA (with the Geisser-Greenhouse correction) and post-hoc Tukey’s multiple comparisons test with individual variances computed for each comparison. (D) Serum-starved control or dynamin TKO MEFs were stimulated with PDGF-BB (50 ng/ml) for 10 min. Lysates were subjected to immunoblotting with indicated antibodies. Representative data are shown from one of 2 independent experiments. (E) Serum-starved Swiss 3T3 fibroblasts were subjected to glucose uptake assay with PDGF-BB (50 ng/ml), PDGF-BB-conjugated microbeads, or unconjugated PDGF and microbeads in the presence of 2DG for 10 min. Graph shows average 2DG6P level in 3 wells for each condition (technical replicates, n = 3) normalized to total cellular protein. Error bars represent ± SD. P values were calculated using ANOVA and post-hoc Tukey’s tests with unstimulated cells set as control. Representative data are shown from one of 2 independent experiments. (F) Serum-starved Swiss 3T3 fibroblasts were pre-treated with dynein inhibitors Ciliobrevin D (30 μM) or Dynarrestin (30 μM) for 20 min or left untreated and then unstimulated or stimulated with PDGF-BB (50 ng/ml) in the presence of 2DG for 10 min. Graph shows average 2DG6P formation in 3 wells for each condition (technical replicates, n = 3) normalized to total cellular protein. Error bars represent ± SD. P values were calculated using ANOVA and post-hoc Tukey’s tests. Representative data are shown from one of 2 independent experiments. (G) Swiss 3T3 fibroblasts ectopically expressing mCherry-fused PDGFR (magenta) with or without Green Glifon4000 glucose sensor or EGFP (green) were serum-starved and stimulated with PDGF-BB (50 ng/ml) for 10 min. Higher magnification images of the boxed regions are shown on the right. Representative data are shown from one of 5 independent live cell imaging experiments. Scale bar: 20 μm. (H) Serum-starved Swiss 3T3 fibroblasts were stimulated with PDGF-BB (50 ng/ml), EGF (50 ng/ml), bFGF (50 ng/ml), HGF (50 ng/ml), or insulin (50 ng/ml) in the presence of 2DG for 10 min. Graph shows average 2DG6P levels in 3 wells for each condition (technical replicates, n = 3) normalized to cellular protein. Error bars represent ± SD. P values were calculated by comparing to respective unstimulated cells using ANOVA and post-hoc Tukey’s tests. Representative data are shown from one of 2 independent experiments. (I) Post-nuclear supernatants and endocytic vesicle fraction with or without 1% Triton X-100 were subjected to glycolysis assay. Graph shows lactate production normalized to protein levels in each fraction. Representative data are shown from one of 2 independent experiments.

PDGFR is internalized upon ligand stimulation by clathrin-dependent and, depending on the PDGF dose, clathrin-independent endocytosis (8). Our endocytic vesicle enrichment experiments detected clathrin heavy and light chain as well as caveolin 1 and flotillins (Fig S1E, Table S1), suggesting involvement of clathrin-dependent and -independent processes. Both endocytosis pathways require dynamin, the GTPase that pinches off the invaginated membrane to generate intracellular vesicles (24). There are three dynamin isoforms, with dynamin 2 being the major isoform in fibroblasts (24, 25). To investigate the potential role of GLUT1 endocytosis in glucose uptake, we employed dynamin 1, 2, 3 conditional triple-knockout mouse embryonic fibroblasts (MEFs), which express Cre recombinase fused with estrogen receptor (ER-Cre) (25). As expected, treatment of these cells with 4-hydroxytamoxifen (4-OHT) resulted in inducible dynamin deletion, and as expected, dynamin triple-knockout (TKO) MEFs (unlike controls) failed to show PDGF-dependent endocytosis of PDGFRα (Fig. 3B). Notably, PDGF-stimulated GLUT1 endocytosis also failed to occur in dynamin TKO MEFs (Fig. 3B). Basal glucose uptake in TKO MEFs was slightly enhanced compared with that in control MEFs (Fig. 3C) and was mediated by GLUT1 in both cells, as indicated by its BAY876 dependence (Fig. S4E). By contrast, while PDGF-BB-stimulated GLUT1-dependent glucose uptake in control MEFs, PDGF failed to stimulate glucose uptake in TKO MEFs (Fig. 3C and S4E). These findings indicate that PDGF-evoked glucose uptake requires an intact endocytosis machinery. Compared with controls, TKO MEFs had slightly reduced PDGF-BB-induced PDGFR tyrosine phosphorylation (Fig, 3D), possibly due to failure to inactivate protein-tyrosine phosphatases such as SHP2 via ligand-dependent endosomal ROS generation (26). Nevertheless, these results exclude the possibility that the endocytosis-deficient cells are unable to respond to PDGF. The slightly elevated basal glucose uptake in unstimulated TKO cells compared to unstimulated control cells (Fig. 3C) presumably reflects increased total and/or surface GLUT1 (Figs. 3D, S4F), owing to a lower level of basal GLUT1 endocytosis and degradation.

Pre-treatment of Swiss 3T3 fibroblasts with the MEK inhibitor AZD6244 or the PI3K inhibitor BKM120 efficiently diminished downstream ERK1/2 or AKT phosphorylation but did not interfere with PDGF-BB-dependent PDGFR-GLUT1 co-endocytosis or impair PDGF-stimulated glucose uptake (Fig. S4G-I). These data suggest that the RAS-ERK MAPK and the PI3K/AKT pathways, two major PDGFR-dependent signaling pathways, are not involved in regulating ligand-dependent receptor endocytosis and glucose uptake, and again indicate coincidence between GLUT1 endocytosis and enhanced glucose uptake.

The necessity of PDGFR endocytosis for stimulation-induced glucose uptake was supported further by the observation that PDGF-BB conjugated to 3 μm diameter microbeads did not stimulate glucose uptake of Swiss 3T3 fibroblasts (Fig. 3E). Beads this large cannot be incorporated into the endocytosis pathway, although they induce comparable receptor phosphorylation (Fig. S4J). Importantly, unconjugated microbeads did not impair PDGF-BB-stimulated glucose uptake or PDGR phosphorylation (Fig. 3E, S4J).

The finding that PDGF enhances glucose uptake in a receptor endocytosis-dependent manner, without increasing cell surface GLUT1, suggested increased activity of GLUT1 and/or HKs. A previous report indicated that PKC-stimulated phosphorylation of GLUT1 on S226 activates its transporter function and suggested an involvement of this mechanism in vascular endothelial growth factor (VEGF)-dependent glucose uptake (6). Although PDGF-dependent GLUT1 phosphorylation at S226 was detected by using a phospho-specific antibody, phosphorylation of this site was increased in TKO MEFs, making it is unlikely that S226 phosphorylation alone is responsible for PDGF-enhanced glucose uptake (Fig. 3D). To assess the alternative possibility that PDGF potentiates HK activity independent of glucose supply via transporter(s), we incubated PDGF-stimulated Swiss 3T3 fibroblasts in the presence of 1% Triton X-100, 1 mM 2DG, and 0.5 or 1 mM ATP, and quantified 2DG6P production. Under these conditions, there was no apparent effect of PDGF on HK activity (Fig. S4K).

### Endocytic vesicles act as vehicles for glucose uptake

Taken together, our data demonstrate that PDGF enhances cellular glucose uptake via receptor endocytosis and independent of known mechanisms such as transcriptional, translational, post-transcriptional regulation of pathway component proteins, or PKC-dependent phosphorylation of GLUT1. Conceivably, PDGFR/GLUT1-positive endocytic vesicles could contribute to localized delivery of glucose to mitochondrial HKs; i.e., growth factor stimulation increases HK activity by controlling substrate access, not intrinsic enzyme activity. Endocytic vesicles are transported along microtubules by dyneins (27), and notably, cytoplasmic dynein heavy chain 1 (DYHC1) was enriched in our endocytic vesicle fractionation (Fig. S1E, Table S1). Treatment of Swiss 3T3 fibroblasts with the specific dynein inhibitors ciliobrevin D or dynarrestin (28) did not affect basal glucose update, but effectively suppressed PDGF-dependent enhancement of uptake (Fig. 3F). The observations that dynamin depletion or dynein inhibition did not suppress basal glucose uptake (Fig. 3C, 3F), while BAY876 almost completely suppressed both basal and stimulated glucose uptake (Fig. 3A), indicate the existence of an endocytosis-and vesicle traffic-dependent, active pathway for GLUT1-mediated glucose uptake in addition to an endocytosis-and vesicle traffic-independent, passive mechanism.

To evaluate further the hypothesis that PDGFR-positive endocytic vesicles function as glucose-loaded vehicles, we employed the EGFP-derived glucose sensor Green Glifon4000, which exhibits enhanced fluorescence intensity in the presence of glucose but not of its catabolites such as glucose-6-phosphate (29). As GLUT1 is a facilitated glucose transporter, intra-endosomal glucose could be transported into the cytoplasm, creating local glucose gradients. Consistent with this idea, live cell imaging of PDGF-BB-stimulated Swiss 3T3 fibroblasts ectopically expressing cytoplasmic Green Glifon4000 showed enhanced fluorescence at vesicles near signals from PDGFRα or PDGFRβ fused with mCherry (Fig. 3G).

Stimulation of glucose uptake in MEFs by basic fibroblast growth factor (bFGF)-and hepatocyte growth factor (HGF) also required dynamin, indicating that this mechanism is common to multiple growth factors (Fig. 3H). Notably, insulin did not significantly evoke glucose uptake in these cells (Fig. 3G), possibly reflecting low insulin receptor levels, as it also did not stimulate increased total tyrosine-phosphorylation in these cells (Fig. S4M).

In control MEFs, PLA experiments confirmed that PDGF stimulation resulted in the juxtaposition of PDGFRβ with the glycolytic enzymes HK2, ALDOA, GAPDH, PGK1, ENO1, PKM1/2, or GLUT1, as observed in Swiss 3T3 fibroblasts. By contrast, there was no PDGF-dependent increase in PLA signal between PDGFRβ and glycolytic enzymes except GLUT1 in TKO MEFs (Fig. S5). As these cells differ primarily in their ability to enable growth factor-induced endocytosis, we conclude that PDGF-induced adjacency of glycolytic enzymes and PDGFR requires, and occurs on, receptor-containing endocytic vesicles. In agreement with this conclusion, incubation of nanoparticle-isolated endocytic vesicle fractions in vitro in the presence of glycolytic substrates, glucose, ATP, ADP, and NAD+, produced lactate, the end product of glycolysis, more effectively than an unenriched post nuclear supernatant (Fig. 3I). Furthermore, pre-treatment of the endocytic vesicle fraction with 1% Triton X-100 decreased lactate production (Fig. 3I). Taken together, these data support the notion that glycolytic enzymes form a multi-enzyme complex on receptor-containing endocytic vesicles to efficiently and concertedly execute the reactions of glycolysis.

### Endocytosis machinery is necessary for the cellular carbon metabolism

To investigate the biological significance of these observations, we compared the effects of glucose-limitation on the survival of control and TKO MEFs. Both cell types survived for at least 3 days in standard high glucose DMEM (4.5 g/L glucose) in the presence of 20 ng/ml PDGF-BB without pyruvate and fetal bovine serum (FBS), and both died when glucose was absent from the media (Fig. 4A). By contrast, limiting glucose from 4.5 g/L to 0.2 g/L or 0.1 g/L in the presence of PDGF for 3 days diminished the survival of the endocytosis-deficient TKO cells when compared with control cells (Fig. 4A), signifying that dynamin depletion can induce cell death that is related to cellular glucose metabolism.

**Figure 4.**
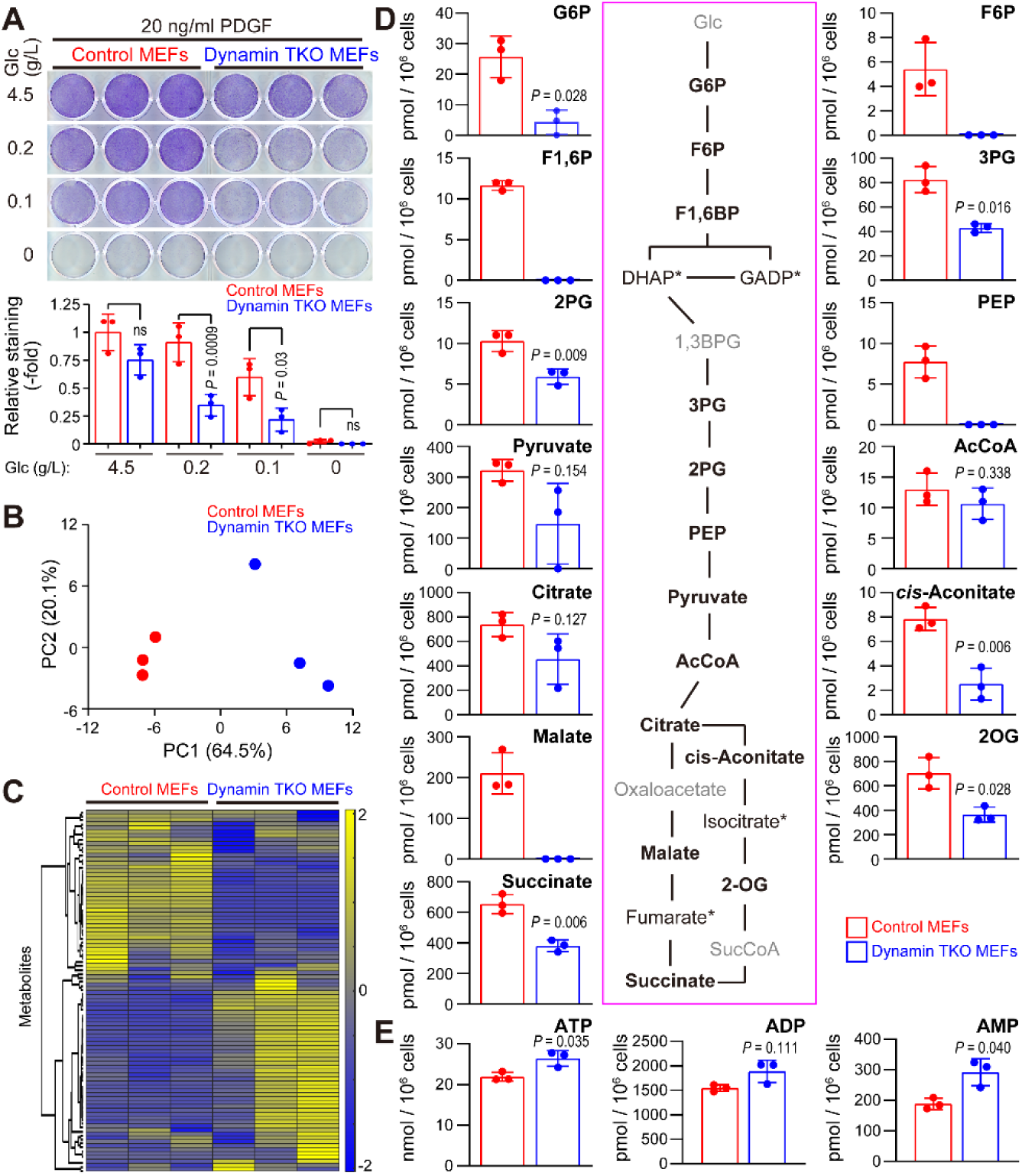
Cellular carbon homeostasis in glucose-limited condition requires receptor endocytosis-dependent glucose uptake. (**A**) Conditional Dynamin TKO MEFs treated with 4-OHT (Dynamin TKO) or untreated (control) were cultured in DMEM containing the indicated concentrations of glucose without FBS and pyruvate in the presence of PDGF-BB (20 ng/ml) for 3 days. Surviving cells were visualized by staining with crystal violet. Graph shows relative staining intensity in 3 wells (technical replicates, n = 3), with the value of control cells set as 1. P values were calculated by ANOVA and post-hoc Tukey’s tests. Representative data are shown from one of 2 independent experiments. (**B**) Control MEFs or dynamin TKO MEFs were cultured in DMEM containing 0.1 g/ml glucose without FBS and pyruvate in the presence of PDGF-BB (20 ng/ml) for 24 h. Cellular metabolites were extracted and analyzed by mass spectrometry. Graph shows principal component analysis. (**C**) Heatmap shows unsupervised hierarchical clustering based on levels of 116 metabolites. (**D**) Graphs show average amount of each metabolite in the 3 samples (technical replicates, n = 3), adjusted by cell numbers. Error bars represent ± SD. P values were calculated using two-tailed Welch’s *t* tests. Scheme shows glycolytic and TCA cycle metabolites. Metabolites not tested are labeled in gray, and asterisks indicate metabolites that were not detected in any sample. (**E**) Graphs show average amount of each metabolite in 3 samples (technical replicates, n = 3), normalized to cell number. Error bars represent ± SD. P values were calculated using two-tailed Welch’s *t* tests. Data was obtained from a single experiment.

To gain further insight into the metabolic consequences of dynamin depletion, we quantified 116 cellular carbon metabolism-related substances in control and TKO MEFs cultured under glucose-limiting conditions (0.1 g/l glucose, without pyruvate) in the presence of PDGF-BB for 24 h, a time at which their viability is comparable. Metabolite levels were normalized to cell number at the time of harvest (Table S2). Principal component analysis and unsupervised hierarchical clustering separated the control and TKO MEF samples (n=3 each) by their genotypes (Fig. 4B, C). Twenty-seven (27) metabolites showed statistically significant alteration between groups, while 13 were detectable only in control or TKO MEF samples (Table S2). Notably, multiple glycolytic intermediates, including glucose-6-phosphate (G6P), fructose-6-phosphate (F6P), fructose-1,6-bisphosphate (F1,6P), diphosphoglycerate (including 1,3-bisphosphoglyceric acid; 1,3BPG), 3-phosphoglyceric acid (3PG), 2-phosphoglyceric acid (2PG), and phosphoenolpyruvic acid (PEP), were strongly diminished in TKO MEFs (Fig. 4D, Table S2). The pentose phosphate pathway (PPP) also was affected dramatically, with 6-phosphogluconic acid (6-PG), ribose 5-phosphate (R5P), sedoheptulose 7-phosphate (S7P), and erythrose 4-phosphate (E4P) significantly decreased or showing a trend towards decrease (Fig. S5A). Pyruvate, acetyl-CoA, and citric acid were nominally, but not statistically diminished, potentially reflecting compensatory amino acid uptake/metabolism. However, the downstream TCA cycle components, cis-aconitate, α-ketoglutarate (2-oxoglutarate; 2OG), succinate, and maleic acid were significantly decreased in TKO MEFs (Fig. 4D, Table S2), indicating compensation is incomplete. Despite depression of these pathways, cellular ATP levels were maintained in TKO MEFs after 24 h of glucose depletion to 0.1 g/L, perhaps due to compensatory ATP synthesis from ADP by adenylate kinases (30), as suggested by the 1.6-fold increase in AMP levels in these cells (Fig. 4E, Table S2). The NADH/NAD^+^-ratio was unaffected, while consistent with the observed decrease in PPP activity, NADPH/NADP^+^ was diminished in the dynamin TKO MEFs (Fig. S5B). Other metabolomic characteristics of dynamin TKO MEFs compared to controls included increased purine metabolic intermediates, unaffected glutathione, nicotinamide, and lipid metabolism, and altered amino acid levels (Table S2, Fig. S5C). Overall, metabolic profiling demonstrated an important role for dynamin, and presumably, dynamin-dependent endocytic events, in central carbon metabolism pathways, including glycolysis, the pentose phosphate pathway, and the TCA cycle. Most likely, these alterations reflect a requirement for dynamin/dynamin-mediated endocytosis in receptor-stimulated glucose uptake, and are essential for survival in limiting glucose.

## Discussion

The current textbook model for glycolysis holds that extracellular glucose molecules are transported through the plasma membrane and then subjected to metabolic reactions in the cytoplasm. Regulation of this process is believed to involve quantitative or qualitative changes in cell surface transporters or glycolytic enzymes. In this study, we found that glucose uptake via GLUT1, heretofore believed to be, with few exceptions, a constitutive, basal glucose transporter, can be enhanced by growth factor action that involves cellular membrane dynamics, particularly endocytosis and vesicle trafficking. Although we cannot exclude the possibility that PDGFR evokes an as yet unknown endocytosis-dependent signaling event that enhances the activity of cell surface GLUT1, we favor a model in which the endocytosis machinery plays a critical role in facilitating re-localization of GLUT1/glycolytic enzymes. We envision two potential mechanisms whereby endocytosis might promote GLUT1-dependent glucose uptake. First, endocytic vesicles could serve as platforms that activate GLUT1 and/or glycolytic enzymes. Although phosphorylation on S226 of GLUT1 per se was insufficient to induce glucose uptake in the context of PDGF stimulation, the signaling enzymes that are enriched on the endocytic vesicles (Fig. S1F, Table S1) could contribute to functional modification of glycolysis-related transporter and/or enzymes. Additionally, endocytosis-dependent protein-protein or protein-lipid interactions on the endosomes might promote transporter/enzyme activity. Notably, direct activation of HK1 on the mitochondria by an isoform of KRAS was reported previously (31). Furthermore, modifications of phosphatidylinositol can occur on endocytic vesicles (32, 33), and we see enrichment of phosphatidylinositol-related enzymes in our vesicle preparation (Fig. S1F, Table S1). Alternatively, induced proximity of glucose-loaded cargos near mitochondria and release of glucose to the adjacent cytoplasmic space might lead to accelerated G6P production simply because of the increase in local glucose concentration, given that basal intracellular glucose is quite low and diffusion in the cytosol could be too slow for efficient activation of glycolysis, perhaps because of molecular crowding (34). The presence of a glycolytic enzyme complex, implied by this study and hypothesized previously to exist in neurons and in muscle (35, 36), might enhance glycolysis by efficiently delivering substrate and concentrating intermediates locally, thereby biasing reactions that are largely at equilibrium under standard conditions.

Growth factors are cues for diverse cellular processes including proliferation, survival, differentiation, morphogenesis, and movement. Immediate enhancement of glucose uptake/glycolysis could prepare cells for functions that likely require a significant amount of energy. The relative importance of ligand-and endocytosis-dependent immediate upregulation and transcription/translation-dependent long-term enhancement of glycolysis may differ depending on the precise cellular context. Notably, our endocytic vesicle enrichment method detected several SLC family transporters including those for amino acids, purines, and ions (Table S1). These findings raise the possibility that endocytic vesicles comprise a third general nutrient intake system using membrane dynamics, in addition to pinocytosis and macropinocytosis (37). Regardless, further studies are required to clarify the involvement of receptor endocytosis in the control of cellular carbon metabolism.

## Materials and Methods

### Antibodies, growth factors, and chemical compounds

Rabbit monoclonal anti-EEA1 (C45B10, #3288), anti-Hexokinase I (C35C4, #2024), anti-GAPDH (D16H11, #5174), anti-PKM 1/2 (C103A3, #3190), anti-PKM2 (D78A4, #4053), and anti-pY849 PDGFRα/pY857PDGFRβ (C43E9, #3170S) antibodies were purchased from Cell Signaling Technology. Rabbit monoclonal anti-GLUT1 (ab115730), anti-Hexokinase II (ab209847), anti-PFKL (ab181064), anti-PGK1 (ab199438), and anti-ENO1/2/3 (ab189891) antibodies were from Abcam. Goat polyclonal anti-PDGFRα antibodies (AF1062) were purchased from R&D systems. Rabbit polyclonal anti-pS226 GLUT1 antibody (ABN991) and mouse anti-phosphotyrosine monoclonal antibody cocktail 4G10 Platinum (05–1050) were purchased from Millipore. Recombinant polyclonal anti-ALDOA antibody cocktail (711764) was purchased from Invitrogen. Goat polyclonal anti-PDGFRβ (sc-1627), anti-EEA1 (sc-6414), anti-clathrin heavy chain (sc-6579), anti-dynamin II (sc-6400), and rabbit polyclonal anti-PDGFRβ (sc-432) antibodies were purchased from Santa Cruz Biotechnology but are currently discontinued. Mouse monoclonal anti-SHP2 (B-1, sc-7384), anti-TPI1 (H-11, sc-166785), anti-GPI (H-10, sc-365066), and anti-PGAM1/4 (D-5, sc-365677) antibodies were purchased from Santa Cruz Biotechnology. Alexa Fluor-conjugated donkey polyclonal anti-goat IgG (ab150130), anti-mouse IgG (ab150105), and anti-rabbit IgG (ab150073) secondary antibodies were purchased from abcam. All antibodies were used at the concentrations recommended by their manufacturers. EGFP-conjugated GLUT1 ligand (receptor binding domain of the human T cell leukemiavirus (HTLV) envelope glycoprotein) was purchased from Metafora Biosystems.

Biotinylated recombinant human PDGF-BB (BT220) and EGF (236-EG) were purchased from R&D Systems. Recombinant human PDGF-BB (500-P47), basic FGF (100-18B), and HGF (100-39H) and were purchased from Peprotech. Recombinant human insulin (099–06473) was purchased from FUJIFILM Wako Chemicals. Streptavidin-coated magnetic iron oxide nanoparticles (SHS-10-05) and MS300/streptavidin Magnosphere microbeads (J-MS-S300S) were purchased from Ocean NanoTech and from JSR Corporation, respectively. 4-hydroxytamoxifen (4-OHT, 17308) was purchased from Cayman Chemical; BAY876 (6199) was purchased from Tocris Bioscience; Selumetinib (AZD6244, S1008) was purchased from Selleck; Buparlisib (BKM120, HY-70063) was purchased from MedChemExpress; ciliobrevin D (250401) and dynarrestin (SML2332) were purchased from Sigma-Aldrich.

### Plasmids

pcDNA3.1/Green Glifon4000 (addgene #126208) was kindly provided by Dr. Kitaguchi (Tokyo Institute of Technology). Expression vectors for mouse PDGFRα and PDGFRβ were generated from pcDNA5FRT-EF-Pdgfra-EGFPN (Addgene #66787) and pcDNA5FRT-EF-Pdgfrb-EGFPN (Addgene #66787), kindly provided by Dr. Pedersen (University of Copenhagen), by replacing EGFP-coding sequences with mCherry-coding sequences using In-Fusion^®^ HD Cloning Kit (Clontech).

### Cell culture and transfection

Swiss 3T3fibroblasts (3T3 Swiss albino) and Dynamin TKO (*Dnm1^fl/fl^*, *Dnm2^fl/fl^*, *Dnm3^fl/fl^*,) MEFs expressing Cre-ER^Tam^ (24) were cultured in Dulbecco-modified Eagle’s medium (DMEM) supplemented with 10% fetal bovine serum (FBS). To induce deletion of dynamin genes, dynamin TKO MEFs were treated with 1 μM 4-OHT for 3 days, followed by additional culture for 7 days before experimentation. Cells without 4-OHT treatment were used as controls. Swiss 3T3 cells were from JCRB Cell Bank (National Institutes of Biomedical Innovation, Health and Nutrition, Japan), and Dynamin TKO MEFs were kindly provided by Dr. De Camilli (Yale University). All cells were confirmed as mycoplasma-negative by the PCR method as reported previously (38).

Swiss 3T3 cells were reverse transfected with the indicated siRNAs using Lipofectamine RNAiMAX (Thermo Fisher), according to the manufacturer’s protocol. Forty-eight hours post-transfection, cells were re-seeded for experiments. Pre-designed siRNAs targeting mouse *Slc2a1*, *Hk1*, *Hk2*, *GPI*, *Aldoa*, *Tpi1*, *Gapdh*, *Pgk1*, *Eno1*, and *Pkm1* were purchased from Sigma-Aldrich.

### Electron microscopy

Swiss 3T3 cells (4 x 10^5^ per dish) were seeded in a 60-mm dish, and then serum-starved for 16 h. Streptavidin-coated iron oxide nanoparticles (final concentration 2 nM) and biotin-PDGF-BB (final concentration 14 nM) were mixed in serum-free DMEM at 37°C for 10 min. Cells were stimulated by incubating in the PDGF/nanoparticle-containing DMEM for the indicated times. Cells were then rinsed with PBS and fixed in fixative containing 2.5% glutaraldehyde and 2% paraformaldehyde at room temperature (RT), then scraped and pelleted into microtubes, and consecutively fixed overnight at 4°C, Cells were post fixed in 1% OsO4, dehydrated in a series of ethanol solutions (30%, 50%, 70%, 85%, 95%, 100%), and embedded in EMbed812 epoxy resin (Electron Microscopy Sciences, Hatfield, PA). 70nm ultrathin sections were cut, mounted on copper grids and stained with uranyl acetate and lead citrate by standard methods. Grids were viewed using a Philips CM12 TEM (Philips) transmission electron microscope and photographed with Gatan 4k x 2.7k digital camera (Gatan Inc.).

### Isolation of PDGFR endocytic vesicles using magnetic nanoparticles

Swiss 3T3 cells (2.5 x 10^6^ per dish) were seeded in 150-mm dishes, and then serum-starved for 16 h. Streptavidin-coated iron oxide nanoparticles (final concentration 2 nM) and biotin-PDGF-BB (final concentration 14 nM) were mixed in serum-free DMEM at 37°C for 10 min. Three 15-mm dishes of cells were stimulated by incubating in the PDGF/nanoparticle-containing DMEM for 5 min. Cells were then rinsed with ice-cold acidic solution (20 mM acetic acid, 150 mM NaCl) once, followed by rinsing with ice-cold PBS twice. After aspirating the PBS, cells were scraped in homogenization buffer (10 mM HEPES pH7.4, 100 mM NaCl, 1 mM EDTA, 10 mM NaF, 10 mM β-glycerophosphate, 2 mM Na3VO4) on ice, centrifuged 800 x *g* for 3 min at 4°C, and re-suspended in 2 ml of homogenization buffer. The plasma membrane was ruptured by nitrogen cavitation or by 30 strokes of Dounce homogenizer (tight pestle). Post-nuclear supernatant (PNS) was prepared by centrifugation of the homogenate 800 x *g* for 5 min at 4°C. A magnetic MS column (Miltenyi Biotec) was set on a magnet and prewashed with 500 μl of homogenization buffer. PNS was applied to the column, followed by washing the column with 3 ml of wash buffer (10 mM HEPES pH7.4, 100 mM NaCl). The column was detached from the magnet and the fraction was eluted with 300 μl of wash buffer by 5 strokes of the syringe plunger.

### LC-MS/MS

Magnetically isolated fractions from Swiss 3T3 fibroblasts treated with PDGF-BB plus unconjugated nanoparticles (control) or PDGF-BB-conjugated nanoparticles (PDGF-particles) were prepared for mass spectrometry analysis as previously described (39). In brief, the samples were reduced with 200mM DTT at 57°C for 1 hour, alkylated with 500 mM iodoacetamide for 45 min at room temperature in the dark, and loaded immediately onto an SDS-PAGE gel and ran just passed the stacking region to remove any detergents and LCMS incompatible reagents. The gel plugs were excised, destained, and subjected to proteolytic digestion using 300 ng of sequencing grade trypsin (Promega) overnight with gentle agitation. The resulting peptides were extracted and desalted as previously described (39).

Aliquots of each sample were loaded onto a trap column (Acclaim® PepMap 100 pre-column, 75 μm × 2 cm, C18, 3 μm, 100 Å, Thermo Scientific) connected to an analytical column (EASY-Spray column, 50 m × 75 μm ID, PepMap RSLC C18, 2 μm, 100 Å, Thermo Scientific) using the autosampler of an Easy nLC 1000 (Thermo Scientific) with solvent A consisting of 2% acetonitrile in 0.5% acetic acid and solvent B consisting of 80% acetonitrile in 0.5% acetic acid. The peptide mixture was gradient eluted into the Orbitrap Fusion Lumos mass spectrometer (Thermo Scientific) using the following gradient: 5%-35% solvent B in 60 min, 35% -45% solvent B in 10 min, followed by 45%-100% solvent B in 10 min. The full scan was acquired with a resolution of 240,000, an AGC target of 1e6, with a maximum ion time of 50ms, and scan range of 400 to 1500 m/z. Following each full MS scan, MS/MS spectra were acquired in the ion trap using the following setting: AGC target of 3e4, maximum ion time of 18ms, one microscan, 2 m/z isolation window and Normalized Collision Energy (NCE) of 32.

All acquired MS2 spectra were searched against a UniProt human database using Sequest within Proteome Discoverer 1.4 (Thermo Fisher Scientific). The search parameters were as follows: precursor mass tolerance ± 10 ppm, fragment mass tolerance ± 0.2 Da, digestion parameters trypsin allowing 2 missed cleavages, fixed modification of carbamidomethyl on cysteine, variable modification of oxidation on methionine, and variable modification of deamidation on glutamine and asparagine. The identifications were first filtered using a 1% peptide and protein FDR cut off searched against a decoy database and only proteins identified by at least two unique peptides were further analyzed. Proteins annotated as common contaminant proteins (cRAP) were excluded and are listed in Table S1. The mass spectrometric raw files are accessible at https://massive.ucsd.edu under accession MassIVE MSV000092208 and at www.proteomexchange.org under accession PXD043112. The following analyses were performed based on data in Table S1.

### Gene annotation enrichment analysis

The list of uniquely detected proteins in the endocytic vesicle fraction were subjected to enrichment analysis to KEGG (40) and Gene Ontology knowledgebases (41) using DAVID Bioinformatics Resources 6.8 (Laboratory of Human Retrovirology and Immunoinformatics (LHRI)) (42). Enriched annotations with the ten lowest P values of each are listed in Fig 1D.

### SDS-PAGE and Immunoblotting

Cells were lysed in SDS lysis buffer (50 mM Tris-HCl pH7.5, 100 mM NaCl, 1 mM EDTA, 1% SDS, 10 mM NaF, 10 mM β-glycerophosphate, 2 mM Na_3_VO_4_). Fractions from the vesicle isolation, or cell lysates were subjected to SDS-PAGE, followed by transfer to Immobilon-P PVDF membranes (Millipore). Membranes were blocked in 1% BSA/TBS containing 0.1% Tween20 for 30 min, and treated with primary antibodies in blocking buffer overnight at 4°C, followed by treatment with HRP-conjugated secondary antibodies (Dako) for 1 h. Bands were visualized using Chemi-Lumi One Super or Chemi-Lumi One Ultra (Nacalai Tesque, 02230-30 and 11644-40) according to the manufacture’s protocol, and images were obtained using an Luminograph I quantitative Cooled CCD camera detection system (ATTO) or an Amersham™ ImageQuant™ 800 (Cytiva) as unsaturated 16-bit TIFF images. Silver staining of gels were performed using Pierce™ Silver Stain for Mass Spectrometry kit (Thermo Fisher, 24600).

### Immunofluorescence

Cells (2 x 10^4^) were seeded on 12 mm, poly-D-lysine-coated circular glass coverslips, and then were serum-starved for 16 h. After stimulation as indicated, cells were rinsed with PBS and fixed in 4% paraformaldehyde/PBS for 10 min at RT, permeabilized with 0.1% Triton X-100, 0.2% BSA/PBS for 10 min, and blocked with 1% BSA/PBS for 30 min, followed by sequential primary antibody and Alexa fluor-conjugated secondary antibody treatments as indicated for 1 h each. For PLA, cells were incubated with primary antibodies, and then treated with donkey anti-goat (Duolink In Situ PLA Probe Anti-Goat MINUS, Sigma) and donkey anti-rabbit (Duolink In Situ PLA Probe Anti-Rabbit PLUS, SIGMA) secondary antibodies, followed by PLA procedure according to the manufacturer’s protocol.

Surface GLUT1 was labeled with a recombinant GLUT1-binding region of human T cell leukemiavirus envelope glycoprotein fused with EGFP. Cells were incubated in the diluted recombinant protein in DMEM (20-fold). To quantify cell surface GLUT1, cells were incubated in the diluted GLUT1-binding protein at 4°C for 30 min and fixed. To observe colocalization of GLUT1 with PDGFR, serum-starved cells were first labeled with the diluted GLUT1-binding protein at 4°C for 30 min, followed by incubation with 50 ng/ml PDGF-BB for 10 min at 37°C, and fixation. Since the recombinant protein showed fluorescence intensity below detectable levels with our equipment on receipt, we visualized the ligand by treating it with an anti-GFP antibody (MBL, #598) and Alexa fluor-conjugated secondary antibody. Coverslips were then mounted on glass slides using Prolongold containing DAPI (Thermo Fisher). Images were obtained using an FV1000 confocal microscopy system (Olympus) as unsaturated 16-bit TIFF images.

### Image analysis

To assess colocalization, regions of interest (ROIs) were set manually according to cell shapes, and Pearson’s correlation coefficients were calculated using the “coloc2” function of FIJI/ImageJ (43) from 8 images per condition.

For PLA images, ROIs were set according to cell shapes and the number of punctate signals generated by PLA were counted using the “Find Maxima” function of FIJI/ImageJ software from 6-10 images per condition.

For surface GLUT1 quantification, total fluorescence intensities, background signal intensities, and cell areas in images were obtained using FIJI/ImageJ software from 6-10 images per condition. Relative fluorescence intensities per cell area were calculated by setting the average intensity of images of control cells to 1.

### Live cell imaging

Swiss 3T3 fibroblasts (5 x 10^4^/10 μl suspension) were transfected with a total 0.5 μg of indicated expression vectors by using a Microporator Mini (Digital Bio Technology, MP-100), setting pulse voltage 1,350 V, pulse width 30 ms, and pulse number 1, according to the manufacturer’s protocol. One hundred thousand (1 x 10^5^) of electroporated cells were then seeded onto a well of a poly-D-lysine-coated µ-Slide 8 Well high Glass Bottom chambered coverslip (ibidi GmbH, 80807). Eight hours after seeding, cells were serum-starved for following 16 h. Cells were observed using an LSM710 confocal microscopy system (Zeiss) with a stage-top incubator at 37°C in phenol red-free, HEPES-buffered DMEM (Nacalai, 09891-25). Images were obtained as unsaturated 16-bit TIFF images.

### Glucose uptake assays

Glucose uptake was quantified using Glucose Uptake-Glo™ Assay kit (Promega) according to the manufacturer’s protocol with some modifications. Cells (2 x 10^4^ per well in 24-well plate or 1 x 10^4^ per well in 48-well plate) were seeded and serum-starved for 16 h. Cells were then rinsed once with DMEM without FBS and glucose, and once with PBS. Cells were then incubated in PBS containing 1 mM 2DG with or without 50 ng/ml of growth factor at 37°C for 10 min. Cellular glucose uptake was terminated by adding acidic Stop Buffer, followed by addition of Neutralization Buffer. The lysates were mixed with the 2DG6P detection reagent, containing G6P dehydrogenase, NAD+, reductase, proluciferin, ATP and recombinant luciferase, and incubated at RT for 1 h. Luminescence signals were quantified using a SpectraMaxiD5 or a SpectraMaxL plate reader (Molecular Devices). Absolute amounts of cellular 2DG6P formation were calculated based on signals of 2DG6P standards, setting the signals of 2DG-untreated cells as the background. 2DG6P formation was normalized by amounts of protein per well. For inhibitor experiments, cells were pre-treated with vehicle or indicated concentrations of BAY876, AZD6244, BKM120, ciliobrevin D, or dynarrestin for 20 min, and then subjected to rinsing and incubation with 2DG in the presence or absence of growth factor. Vehicle or inhibitor was also added to the solutions for rinsing and 2DG/growth factor treatment. For the microbeads experiment, biotinylated PDGF-BB (1.25 μg per well for 24-well plate) was mixed with MS300/streptavidin Magnosphere microbeads (2.0 μg per well for 24-well plate) in 400 μl of PBS and was incubated at RT for 30 min. Beads were then washed 5 times with 1 ml PBS and were resuspended in PBS containing 1 mM 2DG. Cells were treated with the suspension to quantify 2DG6P production.

### In vitro hexokinase assay

Cellular hexokinase activity was measured utilizing the Glucose Uptake-Glo™ Assay kit (Promega) with a modification from the protocol described above. Cells were treated with or without PDGF for 10 min, and rinsed with glucose-free DMEM and PBS. Cells were then incubated in reaction buffer (50 mM Tris–HCl pH 7.4, 10 mM MgCl_2_, 1% Triton X-100, 1 mM 2DG) containing the indicated concentration of ATP, instead of 2DG-containing PBS, at 37°C for 10 min. The reactions were terminated by adding Stop Buffer from the kit, and then the mixtures were neutralized. The mixtures were then mixed with the 2DG6P detection reagent and incubated at RT for 1 h. Luminescence signals were quantified using SpectraMaxiD5 plate reader (Molecular Devices). Absolute amounts of 2DG6P production were calculated based on signals of 2DG6P standards. 2DG6P production was normalized by amounts of protein per well.

### In vitro glycolysis assay

PNS and endocytic vesicle fraction were incubated in the presence or absence of 1% Triton X-100 on ice for 10 min. Protein concentrations of PNS and endocytic vesicle fraction were quantified using DC Protein Assay kit (BIO-RAD) and CBQCA Protein Quantitation Kit (Invitrogen), respectively. Glycolysis reaction was initiated by adding 1/20 volume of PNS or the endocytic vesicle fraction to the reaction mixture (final concentration, 100 mM phosphate buffer pH 7.4, 5 mM NaCl, 2.5 mM MgCl_2_, 0.1 mM DTT, 15 mM ADP, 1 mM ATP, 7 mM NAD^+^, 50 mM glucose). After the indicated time periods of incubation at 37°C, the amounts of lactate in aliquots were quantified immediately using Amplite Fluorometric L-Lactate Assay Kit (AAT Bioquest). Fluorescence signals were quantified using a SpectraMaxiD5 plate reader (Molecular Devices). Reaction mixtures without PNS or endocytic fraction were set as backgrounds. Absolute amounts of lactate were calculated based on signals of standards, and relative lactate productions were calculated by normalizing the values with amounts of protein.

### Cell survival assay

Control or dynamin TKO fibroblasts (1 x 10^5^ per well) suspended in DMEM supplemented with 10% FBS were seeded on 12-well plate and incubated at 37°C in CO_2_ incubator for 16 h. Media were then changed to DMEM without FBS and pyruvate containing PDGF-BB (20 ng/ml) and indicated concentrations of glucose (day 0). Cells were incubated for 3 days, and media were changed every day. On day 3, cells were rinsed with PBS, followed by staining with 0.5% crystal violet solution containing 25% methanol for 30 min. Culture plates were rinsed with tap water and air dried. Plates were then scanned as 16-bit TIFF images, and staining intensities of each well were quantified using ImageJ/FIJI software.

### Metabolomic analysis

Control or dynamin TKO fibroblasts (1.5 x 10^6^), suspended in DMEM supplemented with 10% FBS, were seeded on four 100-mm dishes and incubated at 37°C in a CO_2_ incubator for 16 h. Media were then changed to DMEM without FBS and pyruvate containing 0.1 g/L glucose. Three of the four dishes were used for metabolite extraction, and the other was used for cell counting. Twenty-four hours after medium change, cells were washed twice with 5% mannitol solution, and then treated with 800 μl of methanol and left for 30 s. The cell extracts were then mixed with 550 μl of Milli-Q water containing internal standards (H3304-1002, Human Metabolome Technologies (HMT), Tsuruoka, Yamagata, Japan) and left for another 30 s. The extracts were centrifuged at 2,300 × g and 4°C for 5 min and 800 μl of the aqueous layer was centrifugally filtered through a Millipore 5-kDa cutoff filter (UltrafreeMC-PLHCC, HMT) to remove macromolecules (9,100 ×g, 4°C, 120 min). The filtrate was centrifugally concentrated and re-suspended in 50 μl of Milli-Q water.

The metabolome analysis was conducted with a C-SCOPE package of HMT using capillary electrophoresis time-of-flight mass spectrometry (CE-TOFMS) for cation analysis and CE-tandem mass spectrometry (CE-MS/MS) for anion analysis. Briefly, CE-TOFMS analysis was carried out using an Agilent CE capillary electrophoresis system equipped with an Agilent 6210 time-of-flight mass spectrometer (Agilent Technologies, Waldbronn, Germany). The systems were controlled by Agilent G2201AA ChemStation software version B.03.01 for CE (Agilent Technologies) and connected by a fused silica capillary (50 μm i.d. × 80 cm total length) with commercial electrophoresis buffer (H3301-1001 and I3302-1023 for cation and anion analyses, respectively, HMT) as the electrolyte. The spectrometer was scanned from m/z 50 to 1,000 (44). Peaks were extracted using MasterHands, automatic integration software (Keio University, Tsuruoka, Yamagata, Japan) (45) and MassHunter Quantitative Analysis B.04.00 (Agilent Technologies) in order to obtain peak information including m/z, peak area, and migration time (MT). Signal peaks were annotated according to the HMT metabolite database based on their m/z values with the MTs. The peak area of each metabolite was first normalized with respect to the area of the internal standard and metabolite concentration was evaluated by standard curves with three-point calibrations using each standard compound. Hierarchical cluster analysis (HCA) and principal component analysis (PCA) were performed by HMT’s proprietary software, PeakStat and SampleStat, respectively.

### Statistics and Reproducibility

No statistical method was used to predetermine sample sizes. Samples were not randomized. The investigators were not blinded to allocation during experiments or outcome assessment. Sample sizes and statistical tests for each experiment are denoted in the figure or legends. Each experiment was performed at least twice per condition of the experiment, and representative images from one of the biological replicates are shown in each panel.

Statistical analysis was performed by using the two-tailed Welch’s t test (Figs. 2C, 2E, 4D, 4E, S6A, S6B, S6C, Table S2), unpaired/paired two-tailed t test (Fig. S2B, S4A, S4C), one-way ANOVA with post-hoc Tukey’s test (Figs. 3A, 3E, 3F, 3H, 4A, S4E, S4F, S4H, S4K), RM one-way ANOVA (with the Geisser-Greenhouse correction) and post-hoc Tukey’s multiple comparisons test with individual variances computed for each comparison (Fig 3C), Brown-Forsythe and Welch ANOVA test with post-hoc Games-Howell test (Figs. S3C, S5) by using Graphpad Prism 8 software, where appropriate. Precise P values can be found in the figures.

## Data Availability

Source data for figures are available from the corresponding author upon request.

## Acknowledgments

We thank Dr. P. De Camilli (Yale School of Medicine) for dynamin TKO MEFs. We thank Drs. T. Kitaguchi (Tokyo Institute of Technology) and L. B. Pedersen (University of Copenhagen) for plasmids. We also thank Drs. T. Akaike (Tohoku University), R. L. Possemato (NYU Grossman School of Medicine), and M.R. Philips (NYU Grossman School of Medicine) for helpful comments and discussion. This work was supported by Japan Society for the Promotion of Science (JSPS) 18H06120, 20K06633 (to R.T.), 20H00488 (to Y.S.), and, while R.T. was in the Neel laboratory, by NIH R01 CA49152 (to B.G.N.). R.T. was also supported by the Takeda Science Foundation and the Kato Memorial Bioscience Foundation. We thank NYU Langone Health DART Microscopy Lab for their assistance with TEM work. Microscopy and Proteomics labs are partially funded by NYU Cancer Center Support Grant NIH/NCI P30CA016087. The mass spectrometric experiments were supported with a shared instrumentation grant from the NIH, 1S10OD010582-01A1 for the purchase of an Orbitrap Fusion Lumos.

