## Supplementary Figures S1-S8 for "Endocytic vesicles act as vehicles for glucose uptake in response to growth factor stimulation"

**This PDF file includes:**

Figures S1 to S8

**Other supporting materials for this manuscript include the following:**

Tables S1 and S2

Sup Fig. 1

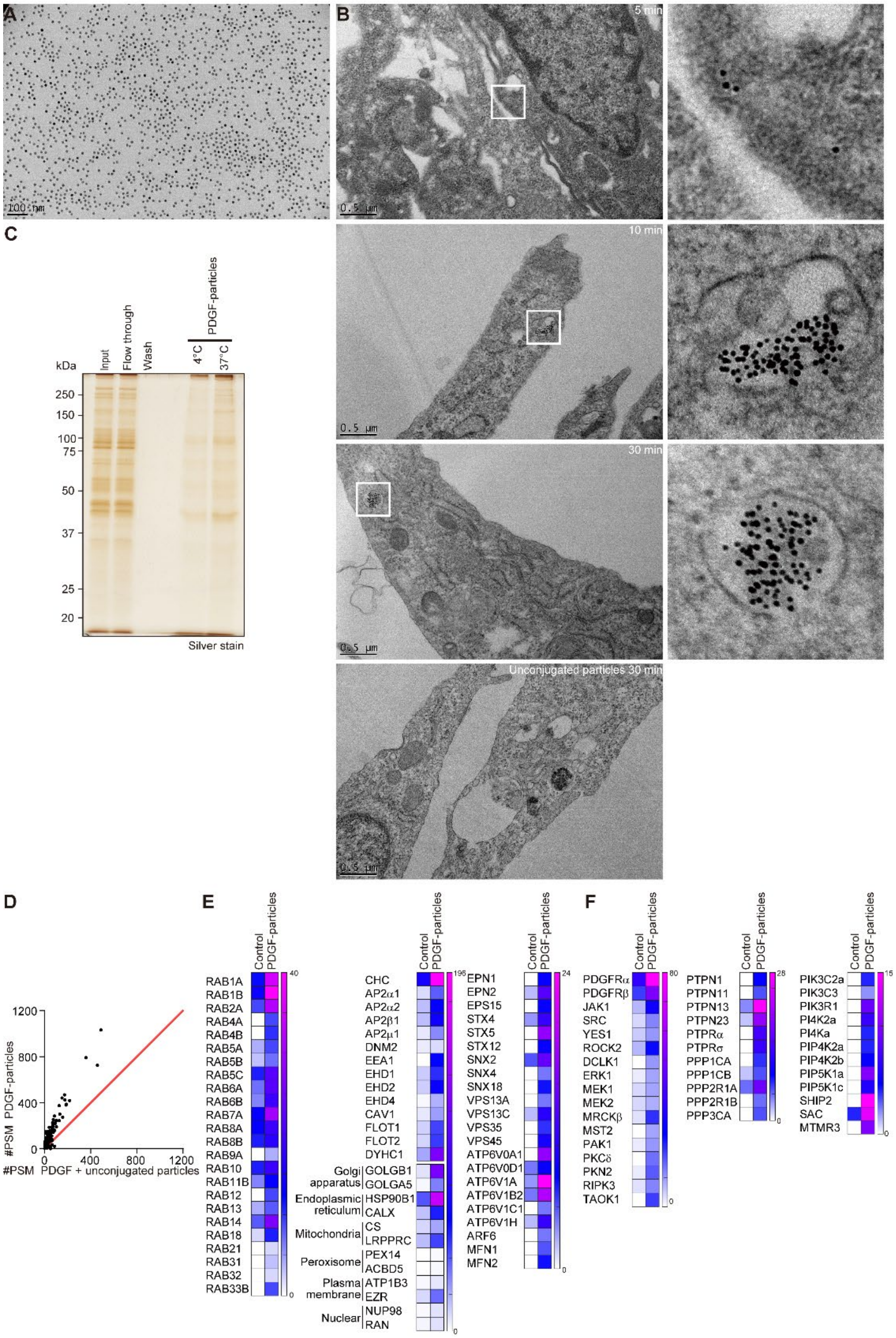


**Fig. S1.**

**Isolation of PDGFR endocytic vesicles utilizing magnetic nanoparticles.** (A) Electron microscopic image of the nanoparticles used in this research. Scale bar: 100 nm. Data were obtained from a single experiment. (B) Serum-starved Swiss 3T3 fibroblasts were treated with PDGF-BB-conjugated or unconjugated nanoparticles for the indicated times. Cells were fixed and analyzed by transmission electron microscopy. Higher magnification images of the boxed region are shown on the right. Scale bars: 0.5 μm. Data was obtained from a single experiment. (C) Serum-starved Swiss 3T3 cells were treated with PDGF-BB-conjugated nanoparticles for 5 min at 4°C or 37°C. Post-nuclear supernatants were prepared and subjected to magnetic isolation, SDS-PAGE, and silver staining. Representative data from one of 2 independent experiments are shown. (D-F) Fractions magnetically isolated from post-nuclear supernatants of PDGF-BB-biotin-conjugated nanoparticle-treated (PDGF-particle) or unconjugated PDGF-BB plus nanoparticle-treated (control) Swiss 3T3 fibroblasts were analyzed by LC-MS/MS. (D) Proteins identified by LC-MS/MS (see also Table S1) were plotted according to #PSM in the control and the PDGF-particle endocytic vesicle fractions. The red line indicates a 1:1 ratio between the fractions. (E, F) Heat maps showing #PSMs of the indicated proteins in PDGF plus unconjugated particles (control) and PDGF-particle endocytic vesicle fractions. Data were obtained from a single experiment.

Sup Fig. 2

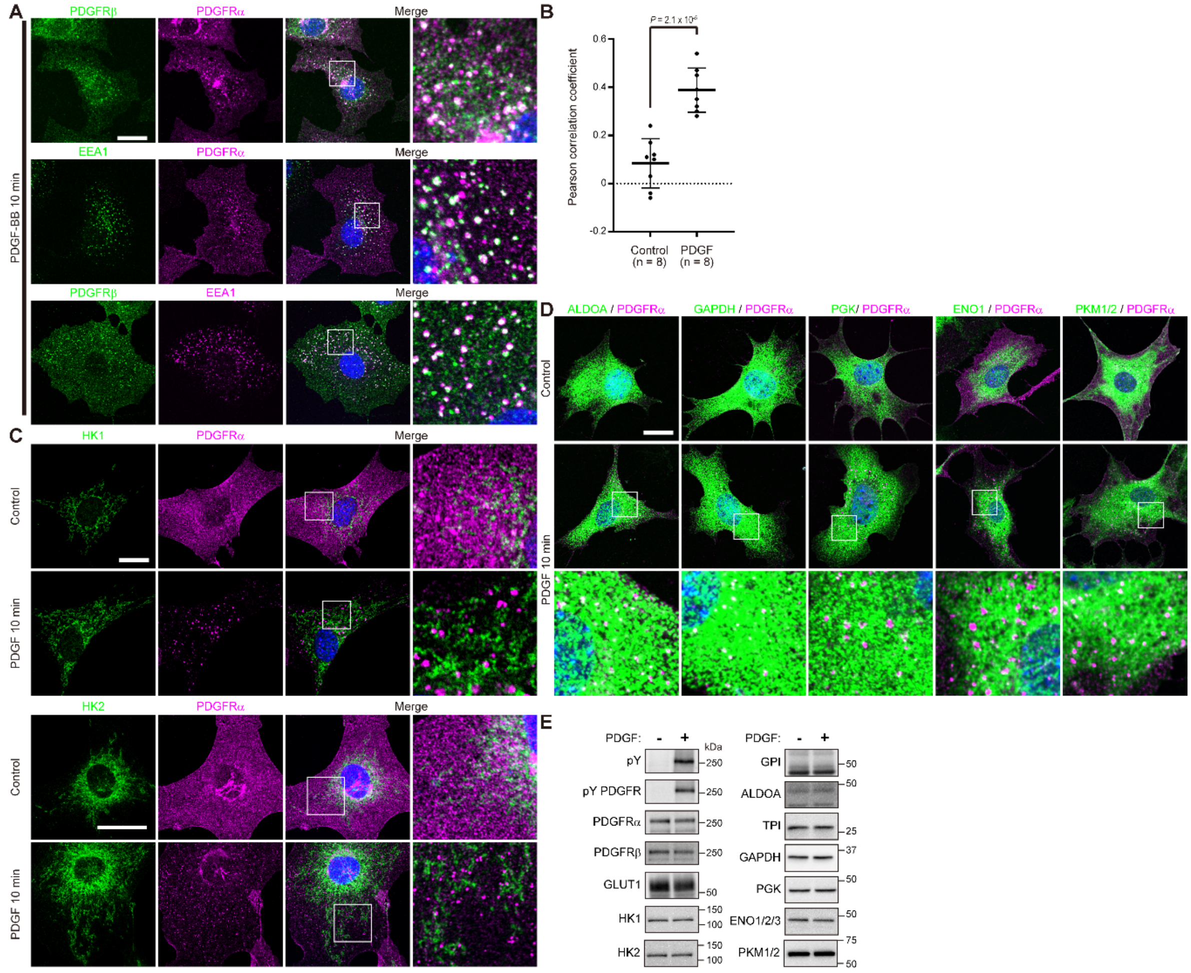


**Fig. S2.**

**Subcellular localization of GLUT1 and glycolytic enzymes in PDGF-treated cells.** (A) Serum-starved Swiss 3T3 fibroblasts were stimulated with 50 ng/ml PDGF-BB for 10 min and subjected to immunofluorescence staining with the indicated antibodies. Higher magnification images of the boxed region are shown on the right. Scale bars: 20 μm. Representative data are shown from one of 2 independent experiments. (B) PDGF-dependent colocalization of PDGFRα and GLUT1 shown in Fig. 2A was quantified. Pearson correlation coefficients were calculated (n = 8 images) and plotted in the graph. Bars represent mean ± SD. P value was calculated using two-tailed unpaired t test. Data are from a single experiment. (C, D) Serum-starved Swiss 3T3 fibroblasts were stimulated with 50 ng/ml PDGF-BB for 10 min and subjected to immunofluorescence staining with the indicated antibodies. Higher magnification images of the boxed region are shown on the right. Scale bars: 20 μm. Representative data are shown from one of 2 independent experiments. (E) Serum-starved Swiss 3T3 fibroblasts were stimulated with or without 50 ng/ml PDGF-BB for 10 min. Lysates were subjected to immunoblotting with the indicated antibodies. Representative data are shown from one of 2 independent experiments.

Sup Fig. 3

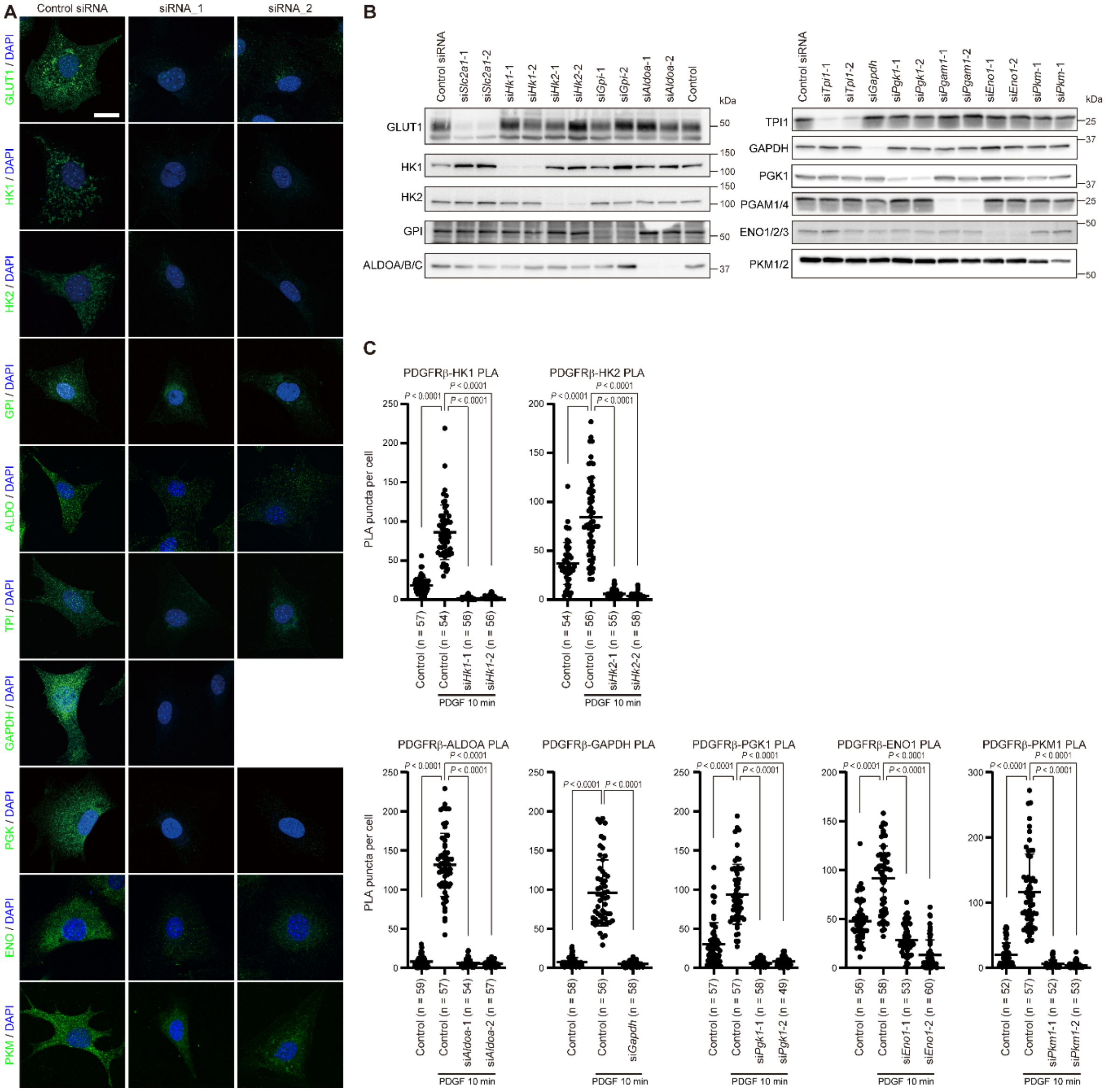


**Fig. S3.**

**Validation of antibodies used in this study and quantification of localization experiments.** (A) Swiss 3T3 fibroblasts were transfected with the indicated siRNAs and cultured for 72 h. Cells were then immunostained with the indicated antibodies (green). Nuclei were stained with DAPI (blue). Scale bar: 20 μm. Representative data are shown from one of 2 independent experiments. (B) Lysates from siRNA-transfected Swiss 3T3 fibroblasts were subjected to immunoblotting with the indicated antibodies. Representative data are shown from one of 2 independent experiments. (C) Serum-starved Swiss 3T3 fibroblasts transfected with indicated siRNAs were stimulated with 50 ng/ml PDGF-BB for 10 min and then were subjected to PLA with anti-PDGFRβ and indicated antibodies. PLA signals in the indicated number of cells were counted and plotted in the graphs. Bars and error bars in the graphs represent mean and ± SD of PLA signals per cell. P values were calculated using Brown-Forsythe and Welch

ANOVA test with post-hoc Games-Howell test. Representative data are shown from one of 2 independent experiments.

Sup Fig. 4

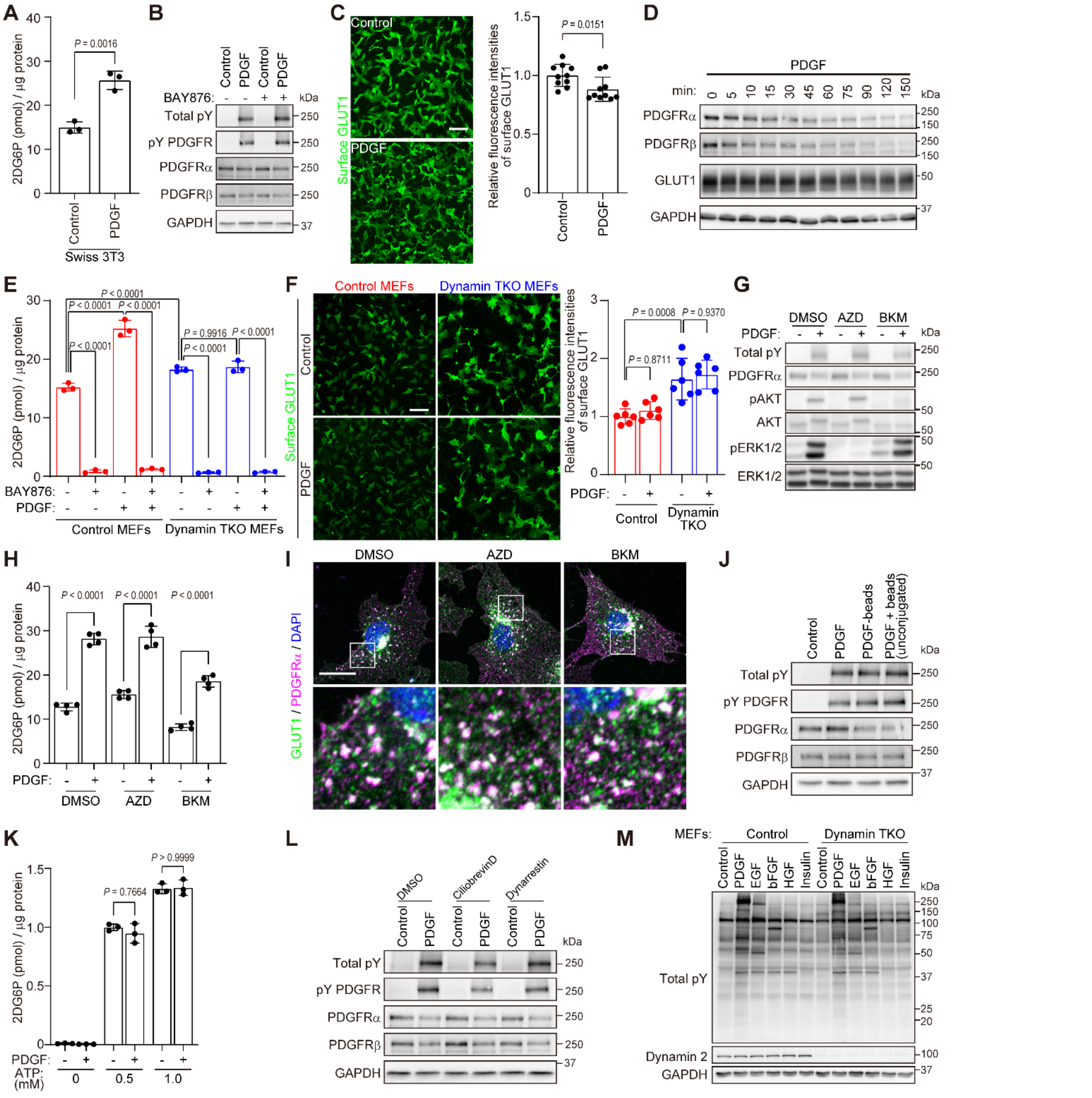


**Fig. S4.**

**Requirement for receptor endocytosis in growth factor-evoked glycolysis.** (A) Serum-starved Swiss 3T3 fibroblasts were subjected to glucose uptake assay in the presence or absence of 50 ng/ml PDGF-BB. Graph shows average 2DG6P level in 3 wells for each condition (technical replicates, n = 3) normalized to total cellular protein. Error bars represent ± SD. P values were calculated using unpaired two-tailed t test. Representative data are shown from one of 2 independent experiments. (B) Serum-starved Swiss 3T3 fibroblasts were pre-treated with BAY876 (50 nM) for 20 min or left untreated, and then were stimulated with PDGF-BB for 10 min or left untreated, as shown in Fig. 3A. Lysates were subjected to immunoblotting with indicated antibodies. Representative data are shown from one of 2 independent experiments. (C) Cell surface GLUT1 was labeled (green) in serum-starved or PDGF-stimulated Swiss 3T3 fibroblasts (as above). Scale bar: 100 μm. Graph shows average fluorescence intensity per cell area in images (10 images each, technical replicates, n = 10), relative to the average in unstimulated cells (normalized to 1). Error bars represent ± SD. P values were calculated by unpaired two-tailed t test. Representative data are shown from one of 2 independent experiments. (D) Serum-starved Swiss 3T3 fibroblasts were stimulated with PDGF-BB (50 ng/ml) for the indicated times. Lysates were subjected to immunoblotting with the indicated antibodies. € Serum-starved untreated (control) or 4-OHT-treated dynamin TKO MEFs were pre-treated with 50 nM BAY876 for 20 min or left untreated, and then were stimulated with PDGF-BB or left unstimulated in the presence of 2DG for 10 min. Graph shows average 2DG6P levels in 3 wells for each condition (technical replicates, n = 3) normalized to cellular protein. Error bars represent ± SD. P values were calculated using ANOVA and post-hoc Tukey's tests. Representative data are shown from one of 2 independent experiments. (F) Serum-starved untreated (control) or 4-OHT-treated dynamin TKO MEFs were stimulated with 50 ng/ml PDGF-BB for 10 min, followed by labeling of cell surface GLUT1 (green) at 4°C for 30 min. Scale bar: 100 μm. The graph shows the average fluorescence intensity per cell area from images (6 images each, technical replicates, n = 6), relative to the average in unstimulated control cells (normalized to 1). Error bars represent ± SD. P values were calculated using one-way ANOVA with post-hoc Tukey's test. Representative data are shown from one of 2 independent experiments. (G-I) Serum-starved Swiss 3T3 fibroblasts were pre-treated with AZD6244 (5 mM) or BKM120 (2.5 mM) for 20 min or left untreated, and then were stimulated with PDGF-BB or left unstimulated. (G) Cells were lysed and subjected to immunoblotting with indicated antibodies. Representative data are shown from one of 2 independent experiments. (H) Cells were then subjected to glucose uptake assay. Graph shows average of 2DG6P in 3 wells for each condition (technical replicates, n = 4), normalized to total cellular protein. Error bars represent ± SD. P values were calculated using ANOVA and post-hoc Tukey's tests. Representative data are shown from one of 2 independent experiments. (I) Cells were subjected to immunostaining with anti-GLUT1 (green) and anti-PDGFRα (magenta) antibodies. Higher magnification images of the boxed regions are shown on the right. Nuclei were stained with DAPI (blue). Representative data are shown from one of 2 independent experiments each. Scale bar: 20 μm. (J) Serum-starved Swiss 3T3 fibroblasts were incubated in the presence or absence of 50 ng/ml PDGF-BB, PDGF-BB conjugated to microbeads, or unconjugated PDGF-BB plus microbeads as shown in Fig. 3F. Lysates were subjected to immunoblotting with the indicated antibodies. Representative data are shown from one of 2 independent experiments. (K) Serum-starved Swiss 3T3 fibroblasts were stimulated with 50 ng/ml PDGF-BB for 10 min. After rinsing with glucose-free DMEM and PBS, cells were incubated in reaction buffer containing 1 mM 2DG, Triton X-100, and the indicated concentrations of ATP for 10 min. Graph shows average 2DG6P levels in 3 wells for each condition (technical replicates, n = 3) normalized to total cellular protein. Error bars represent ± SD. P values were calculated using ANOVA and post-hoc Tukey's tests. Representative data are shown from one of 2 independent experiments. (L) Serum-starved Swiss 3T3 fibroblasts were pre-treated with Ciliobrevin D (30 μM) or Dynarrestin (30 μM) or left untreated for 20 min, and then were stimulated with or without PDGF-BB for 10 min as shown in Fig. 3G. Lysates were subjected to immunoblotting with indicated antibodies. Representative data are shown from one of 2 independent experiments. (M) Serum-starved control or dynamin TKO MEFs were stimulated with 50 ng/ml of the indicated growth factors for 10 min. Lysates were subjected to immunoblotting with the indicated antibodies. Representative data are shown from one of 2 independent experiments.

Sup Fig. 5

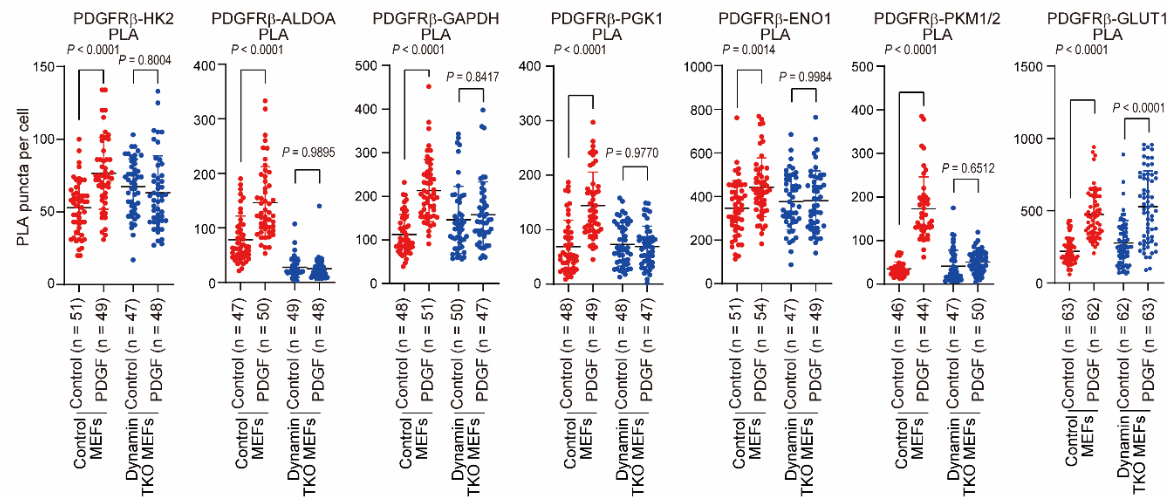


**Fig. S5.**

**Requirement for dynamin in PDGF-dependent clustering of PDGFR and glycolytic enzymes.** Serum-starved untreated (control MEFs) or 4-OHT-treated dynamin TKO MEFs were stimulated with 50 ng/ml PDGF-BB for 10 min, and were then subjected to PLA with anti-PDGFRβ and the indicated antibodies. PLA signals in the indicated number of cells were counted and plotted in the graphs. Bars represent mean ± SD of PLA signals per cell. P values were calculated using Brown-Forsythe and Welch ANOVA test with post-hoc Games-Howell test. Representative data are shown from one of 2 independent experiments.

Sup Fig. 6

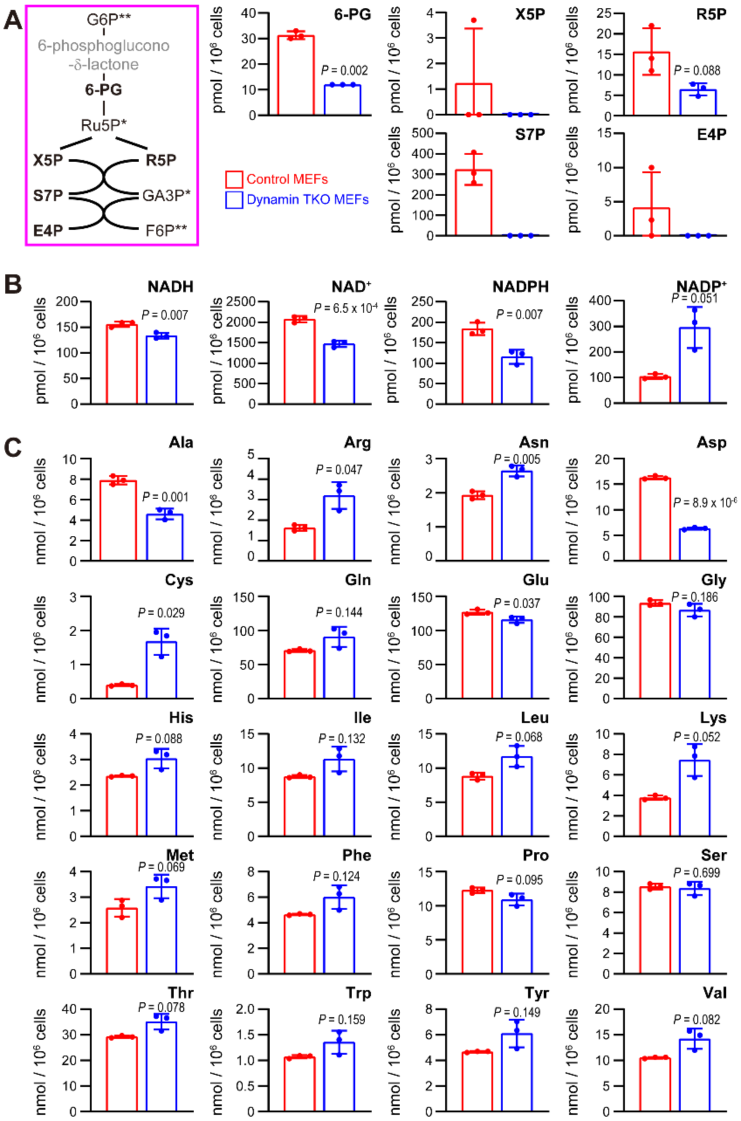


**Fig. S6.**

**Role of receptor endocytosis in cellular metabolism.** (A-C) Graphs show average amounts of each metabolite in 3 samples (technical replicates, n = 3), normalized to cell number. Error bars represent ± SD. P values were calculated using two-tailed Welch's t tests. The scheme in A shows metabolites in the pentose phosphate pathway. Metabolites not tested are labeled in gray; asterisks indicate metabolites that were not detected in any sample. Data were obtained from a single experiment.

Sup Fig. 7

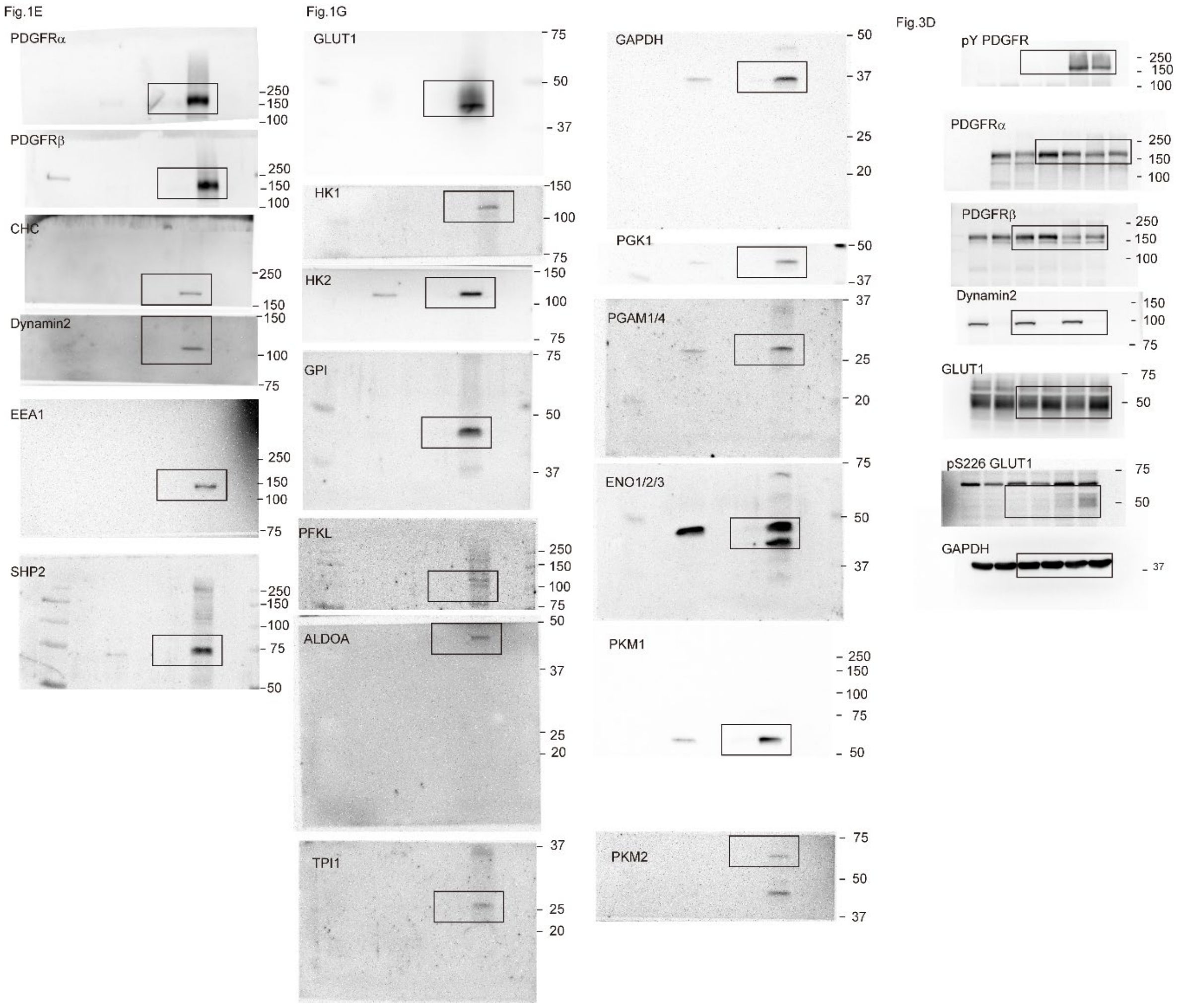


**Fig. S7.**

**Uncropped immunoblot images presented in Figures.**

Sup Fig. 8

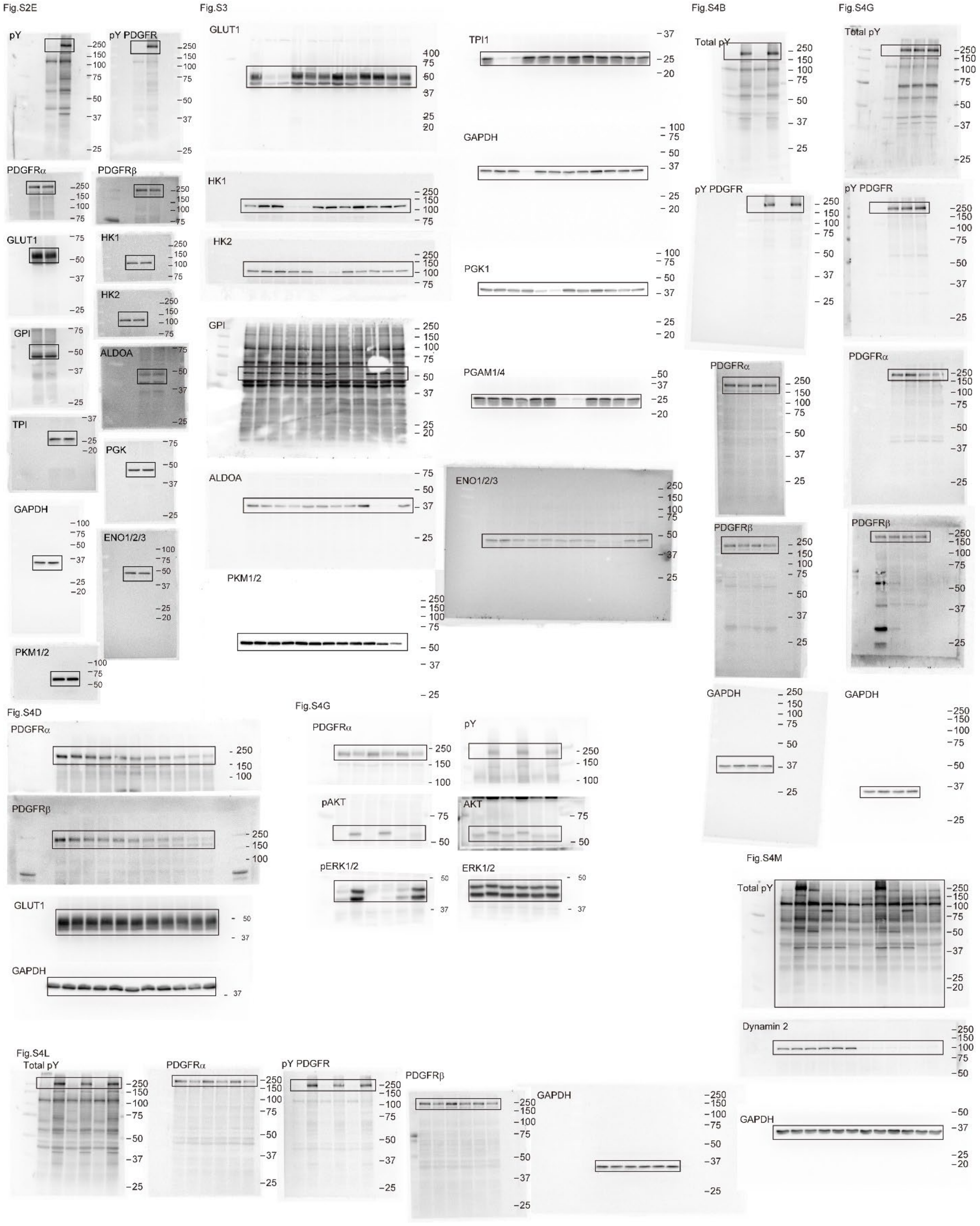


**Fig. S8.**

**Uncropped immunoblot images presented in Figures.**
